## Supplemental data for "Potent but transient immunosuppression of T-cells is a general feature of CD71^+^ erythroid cells"

Tomasz M. Grzywa et al.

**Supplementary Table 1. Clinical data of anemic and non-anemic patients**

| Parameter | | Non-anemic | Anemic | *P* | Reference |
|---|---|---|---|---|---|
| **Sex** | **F** | 23 | 19 | *0.3888* | n/a |
| | **M** | 19 | 23 | | |
| **Age** | | 53.71 ±18.72 | 63.66 ±19.50 | *0.0209* | n/a |
| **WBC** | | 7.12 ±1.76 | 7.59 ±3.44 | *0.9100* | 4.00-11.00 |
| **RBC** | | 4.73 ±0.47 | 3.68 ±0.83 | *<0.0001* | 3.8-5.2 (F)<br>4.2-5.7 (M) |
| **HGB** | | 14.13 ±1.29 | 9.62 ±1.32 | *<0.0001* | 12.0-16.0 (F)<br>14.0-18.0 (M) |
| **HCT** | | 40.99 ±3.87 | 29.73 ±4.10 | *<0.0001* | 37.0-47.0 (F)<br>40.0-54.0 (M) |
| **MCV** | | 86.88 ±3.09 | 83.21 ±14.15 | *0.0371* | 80.0-96.0 |
| **MCH** | | 29.93 ±1.50 | 27.14 ±5.68 | *0.0178* | 27.0-31.2 |
| **RDW-CV** | | 14.88 ±1.752 | 18.90 ±3.54 | *<0.0001* | 11.6-14.8 |
| **%CEC** | | 0.124 ±0.15 | 0.907 ±0.87 | *<0.0001* | n/a |
| **CEC count** | | 4.20 ±5.391 | 26.33 ±28.03 | *<0.0001* | n/a |

**Supplementary Table 2 Individual clinical data of anemic and non-anemic patients**

| Patient | SEX | AGE | WBC | RBC | HGB | HCT | MCV | MCH | RDW-CV | %CECs | CEC count (x $10^3$) |
|---|---|---|---|---|---|---|---|---|---|---|---|
| **H1** | F | 43 | 6,72 | 4,88 | 13,7 | 40,7 | 83,4 | 28,1 | 14,3 | 0,08 | 2,18 |
| **H2** | F | 61 | 6,35 | 4,9 | 14,1 | 41 | 83,7 | 28,8 | 13,4 | 0,13 | 4,11 |
| **H3** | M | 75 | 8,86 | 5,56 | 15,2 | 48,4 | 87,1 | 27,3 | 14,6 | 0,28 | 7,82 |
| **H4** | M | 38 | 7,76 | 4,84 | 14,5 | 42,7 | 88,2 | 30 | 13,2 | 0,16 | 6,35 |
| **H5** | M | 40 | 10,01 | 4,91 | 14,6 | 41,3 | 84,1 | 29,7 | 12,2 | 0,03 | 1,50 |
| **H6** | M | 60 | 7,34 | 4,7 | 14,3 | 40,6 | 86,3 | 30,4 | 16,1 | 0,04 | 1,65 |
| **H7** | M | 29 | 5,07 | 5,02 | 15,2 | 43,3 | 86,2 | 30,2 | 17 | 0,07 | 3,01 |
| **H8** | M | 58 | 4,61 | 4,81 | 14,5 | 40,3 | 87,8 | 30,1 | 17,8 | 0,12 | 5,50 |
| **H9** | M | 64 | 7,49 | 5,04 | 15,6 | 46,2 | 91,7 | 31 | 16,3 | 0,67 | 18,75 |
| **H10** | M | 30 | 6,3 | 5,04 | 15,2 | 44,4 | 88,1 | 30,2 | 14,6 | 0,02 | 1,02 |
| **H11** | F | 53 | 6,92 | 4,53 | 13,1 | 39,3 | 86,8 | 28,9 | 13,6 | 0,24 | 10,48 |
| **H12** | M | 73 | 8,92 | 5,25 | 15,2 | 44,9 | 85,5 | 29 | 15,1 | 0,10 | 5,66 |
| **H13** | F | 60 | 6,02 | 4,65 | 13,5 | 39,7 | 85,4 | 29 | 12,4 | 0,10 | 4,23 |
| **H14** | M | 16 | 9,14 | 5,94 | 16,6 | 48,8 | 82,2 | 27,9 | 14 | 0,17 | 6,34 |
| **H15** | M | 38 | 5,66 | 4,71 | 15 | 41,9 | 88,9 | 31,9 | 15,5 | 0,04 | 1,92 |
| **H16** | M | 42 | 5,14 | 5,64 | 16,5 | 48 | 85,1 | 29,3 | 12,6 | 0,08 | 3,58 |
| **H17** | F | 84 | 10,39 | 4,4 | 13,1 | 37,2 | 84,5 | 29,8 | 12,7 | 0,07 | 2,52 |
| **H18** | M | 23 | 5,1 | 4,88 | 14,7 | 42,8 | 87,7 | 30,1 | 11 | 0,02 | 1,04 |
| **H19** | F | 78 | 6,29 | 4,26 | 13,2 | 38,4 | 90,1 | 31 | 13,3 | 0,10 | 2,64 |
| **H20** | F | 22 | 5,02 | 4,91 | 14,7 | 41,2 | 83,9 | 30 | 16,2 | 0,06 | 1,93 |
| **H21** | F | 55 | 6,33 | 4,13 | 12,7 | 37 | 89,5 | 30,9 | 17,5 | 0,04 | 0,98 |
| **H22** | F | 71 | 10,6 | 4,87 | 15,5 | 44,6 | 91,6 | 31,9 | 16,5 | 0,07 | 2,28 |
| **H23** | F | 42 | 6,41 | 4,29 | 13,2 | 36,8 | 85,8 | 30,8 | 14,9 | 0,03 | 0,89 |
| **H24** | F | 67 | 6,18 | 4,35 | 13,2 | 36,6 | 84 | 30,2 | 15,5 | 0,05 | 1,33 |
| **H25** | F | 70 | 7,45 | 4,14 | 12,1 | 34,6 | 85,3 | 29,3 | 16,7 | 0,52 | 9,85 |
| **H26** | F | 45 | 7,87 | 4,51 | 12,1 | 38,3 | 84,9 | 26,8 | 13,9 | 0,17 | 5,38 |
| **H27** | F | 58 | 7,11 | 3,9 | 12 | 33,5 | 85,8 | 30,9 | 15,3 | 0,28 | 6,93 |
| **H28** | M | 64 | 4,9 | 4,92 | 15,2 | 41,9 | 85,3 | 30,8 | 15,8 | 0,04 | 1,23 |
| **H29** | M | 63 | 9,55 | 5,14 | 14,6 | 44,5 | 86,6 | 28,4 | 14,2 | 0,08 | 2,96 |
| **H30** | M | 28 | 5,57 | 4,98 | 14,7 | 44,8 | 90 | 29,5 | 13,2 | 0,02 | 0,94 |
| **H31** | F | 28 | 6,06 | 4,66 | 12,4 | 38,6 | 82,8 | 26,6 | 15 | 0,04 | 1,51 |
| **H32** | M | 35 | 5,91 | 5,65 | 17,2 | 46,6 | 82,4 | 30,4 | 14,7 | 0,03 | 1,38 |
| **H33** | F | 87 | 9,2 | 4,1 | 12,8 | 36,3 | 88,5 | 31,1 | 18,6 | 0,01 | 0,47 |
| **H34** | F | 45 | 6,95 | 4,31 | 13 | 36,5 | 84,8 | 30,1 | 15,6 | 0,01 | 0,27 |
| **H35** | M | 58 | 10,94 | 4,2 | 14,3 | 40,2 | 95,8 | 34,1 | 16,2 | 0,05 | 1,19 |
| **H36** | F | 80 | 9,82 | 4,27 | 14,1 | 40,4 | 94,6 | 33 | 18,2 | 0,36 | 8,47 |
| **H37** | F | 66 | 5,99 | 4,49 | 12,8 | 40,2 | 89,5 | 28,5 | 15 | 0,10 | 2,38 |
| **H38** | F | 85 | 5,2 | 4,17 | 12,5 | 36,8 | 88,2 | 30 | 12,8 | 0,03 | 0,63 |
| **H39** | F | 49 | 7,22 | 4,2 | 13,1 | 37 | 88,2 | 31,2 | 16 | 0,10 | 2,00 |
| **H40** | M | 61 | 7,94 | 5,17 | 15,2 | 45,6 | 88,2 | 29,4 | 13 | 0,05 | 1,75 |
| **H41** | F | 58 | 5,45 | 4,61 | 14,2 | 38,5 | 83,6 | 30,7 | 15,6 | 0,06 | 1,50 |
| **H42** | F | 58 | 5,45 | 4,61 | 14,2 | 38,5 | 83,6 | 30,7 | 15,6 | 0,51 | 29,85 |
| **A1** | M | 27 | 4,99 | 4,66 | 9,9 | 33,2 | 71,2 | 21,2 | 19 | 0,36 | 9,30 |
| **A2** | M | 41 | 8,64 | 3,69 | 9,2 | 26,8 | 72,7 | 24,8 | 25,3 | 0,64 | 18,46 |
| **A3** | F | 19 | 4,94 | 3,74 | 8,7 | 27,8 | 74,3 | 23,3 | 17 | 0,14 | 1,74 |
| **A4** | F | 51 | 7,79 | 4,74 | 9,1 | 31,3 | 66 | 19,2 | 24,3 | 0,96 | 26,34 |

| | | | | | | | | | | | |
|---|---|---|---|---|---|---|---|---|---|---|---|
| **A5** | M | 71 | 10,66 | 4 | 9,5 | 28,5 | 71,2 | 23,7 | 20,3 | 2,18 | 116,85 |
| **A6** | M | 64 | 5,07 | 5,26 | 11,4 | 37,2 | 70,7 | 21,7 | 19,9 | 0,92 | 48,52 |
| **A7** | F | 44 | 6,64 | 4,17 | 8,9 | 29,1 | 69,8 | 21,3 | 19,5 | 0,09 | 3,08 |
| **A8** | M | 88 | 11,03 | 4,49 | 9,7 | 32 | 71,3 | 21,6 | 25,8 | 0,98 | 30,89 |
| **A9** | F | 63 | 3,26 | 2,25 | 8,8 | 25,3 | 112,4 | 39,1 | 18,7 | 0,32 | 6,05 |
| **A10** | F | 90 | 6,67 | 3,96 | 9,3 | 28,4 | 71,7 | 23,5 | 22,3 | 0,21 | 7,37 |
| **A11** | M | 83 | 8,81 | 3,22 | 10,7 | 32,1 | 99,7 | 33,2 | 15,5 | 2,03 | 56,17 |
| **A12** | F | 23 | 4,01 | 4,3 | 8,4 | 29,6 | 68,8 | 19,5 | 18,1 | 1,04 | 37,98 |
| **A13** | F | 57 | 10,23 | 2,86 | 9,1 | 28 | 97,9 | 31,8 | 12,4 | 0,98 | 17,25 |
| **A14** | F | 44 | 17,24 | 2,63 | 9,1 | 25,7 | 97,7 | 34,6 | 15,5 | 0,47 | 15,30 |
| **A15** | F | 61 | 14,18 | 3,77 | 8,4 | 27,8 | 73,2 | 22,3 | 20,7 | 1,56 | 94,51 |
| **A16** | F | 65 | 5,5 | 4,56 | 8,8 | 29,6 | 64,9 | 19,3 | 25,9 | 1,19 | 67,21 |
| **A17** | M | 64 | 5,99 | 5,07 | 11,1 | 35,9 | 70,8 | 21,9 | 19,2 | 0,68 | 20,62 |
| **A18** | F | 69 | 6,42 | 2,36 | 7,9 | 23,6 | 100 | 33,5 | 19,1 | 3,59 | 45,51 |
| **A19** | M | 66 | 3,41 | 2,51 | 8,5 | 25,4 | 101,2 | 33,9 | 18 | 0,37 | 5,69 |
| **A20** | F | 90 | 4,75 | 4,04 | 9,8 | 28,5 | 70,6 | 24,3 | 22,1 | 0,20 | 5,80 |
| **A21** | F | 70 | 10,53 | 4,4 | 9,7 | 33,1 | 75,2 | 22 | 19,2 | 0,25 | 7,54 |
| **A22** | M | 62 | 9,29 | 3,09 | 9,4 | 29,7 | 96,1 | 30,4 | 15,1 | 1,71 | 56,90 |
| **A23** | M | 59 | 2,92 | 3,18 | 6,5 | 22,8 | 71,8 | 20,4 | 17,2 | 2,45 | 39,86 |
| **A24** | M | 53 | 3,77 | 5,04 | 10,9 | 36,2 | 71,8 | 21,6 | 16,1 | 0,60 | 25,55 |
| **A25** | M | 42 | 5,71 | 2,52 | 8,5 | 25,3 | 100,4 | 33,7 | 18,1 | 0,25 | 6,37 |
| **A26** | M | 57 | 4,88 | 3,07 | 11,3 | 33 | 107,5 | 36,8 | 13,6 | 0,04 | 0,90 |
| **A27** | M | 84 | 6,91 | 2,78 | 9 | 27,4 | 98,6 | 32,4 | 19,8 | 0,28 | 5,94 |
| **A28** | F | 90 | 9,5 | 3,18 | 8,4 | 24,7 | 77,7 | 26,4 | 13,9 | 0,35 | 9,49 |
| **A29** | M | 79 | 11,31 | 3,48 | 8,7 | 24,7 | 71 | 24,9 | 21,4 | 0,12 | 2,62 |
| **A30** | M | 69 | 11,03 | 2,84 | 9,5 | 27,9 | 98,3 | 33,6 | 16,8 | 1,91 | 31,91 |
| **A31** | M | 60 | 6,23 | 3,26 | 11,5 | 34,4 | 105,5 | 35,3 | 13,6 | 1,01 | 80,81 |
| **A32** | F | 86 | 11,86 | 4,33 | 11,9 | 33,5 | 77,4 | 27,5 | 17,4 | 0,01 | 0,45 |
| **A33** | F | 84 | 6,19 | 3,71 | 11,8 | 34 | 91,8 | 31,8 | 18,9 | 0,01 | 0,27 |
| **A34** | F | 90 | 6,68 | 3,98 | 10,5 | 30,9 | 77,6 | 26,4 | 16,4 | 0,09 | 0,73 |
| **A35** | M | 77 | 11,44 | 2,67 | 9,6 | 28,4 | 106,5 | 36 | 19,5 | 2,95 | 23,34 |
| **A36** | M | 77 | 4,5 | 3,06 | 9,5 | 29,4 | 96,1 | 31 | 17,3 | 1,54 | 16,95 |
| **A37** | M | 33 | 10,54 | 4,89 | 13,7 | 40,7 | 83,2 | 28 | 12,7 | 0,09 | 2,27 |
| **A38** | F | 37 | 4,44 | 3,64 | 9,6 | 28,7 | 78,9 | 26,5 | 25,1 | 1,76 | 64,01 |
| **A39** | F | 80 | 3,12 | 4,71 | 10,9 | 36,1 | 76,6 | 23,1 | 23,1 | 1,22 | 38,94 |
| **A40** | M | 60 | 6,04 | 3,36 | 8,5 | 26,4 | 78,6 | 25,3 | 22,3 | 0,12 | 1,72 |
| **A41** | M | 81 | 14,1 | 3,43 | 9 | 25,7 | 75 | 26,1 | 18,9 | 1,53 | 28,13 |

### Supplementary Figures

#### Supplementary Figure 1. Hematological parameters of anemia models

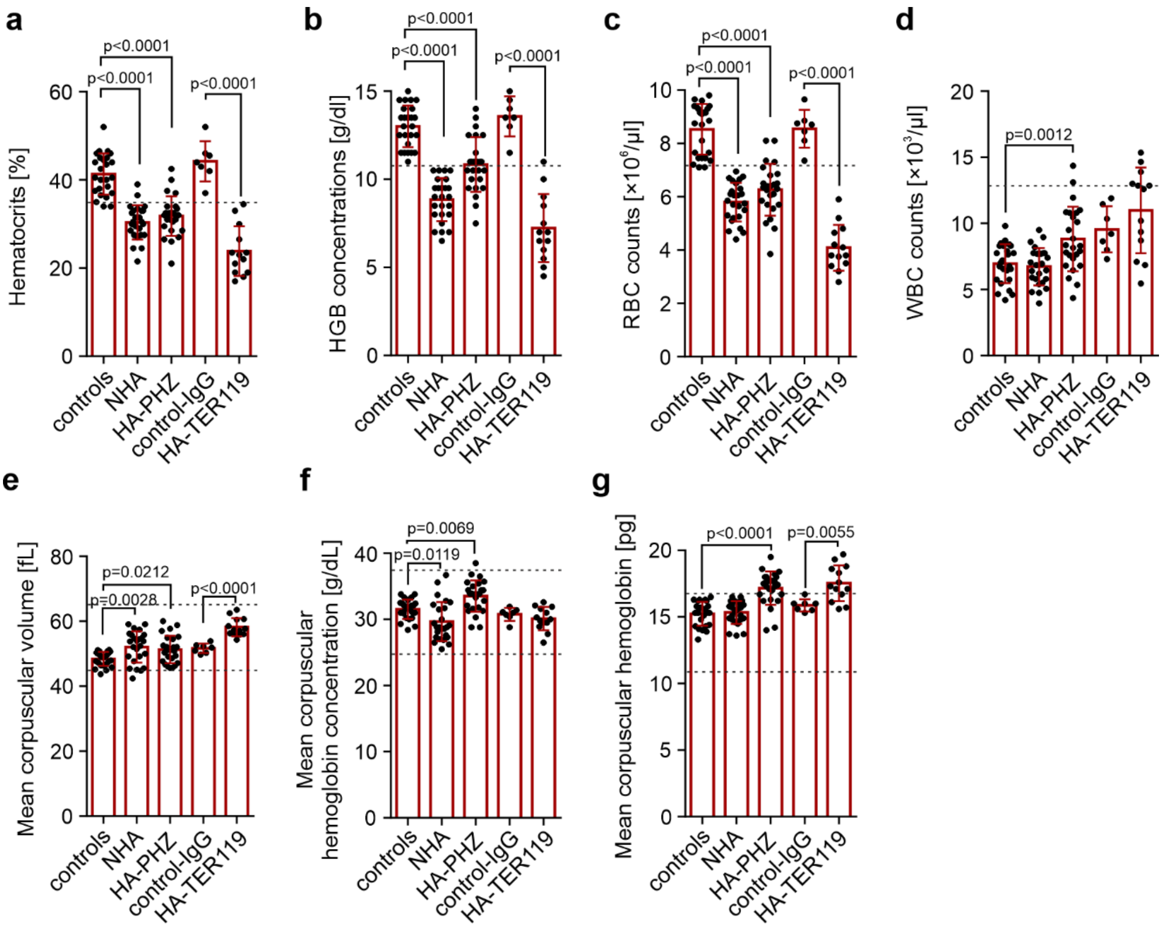


**a-d,** Complete blood count parameters including RBC count (**a**), hematocrit (**b**), hemoglobin concentration (**c**), and WBC count (**d**) in control healthy mice (n=25), control IgG mice (n=7), NHA mice (n=25), HA-PHZ mice (n=25), and HA-TER119 mice (n=13). **e-g,** RBC parameters including mean corpuscular volume (MCV, **e**), mean corpuscular hemoglobin concentration (MCHC, **f**), and mean corpuscular hemoglobin (MCH, **g**) in control healthy mice (n=25), control IgG mice (n=7), NHA mice (n=25), HA-PHZ mice (n=25), and HA-TER119 mice (n=13). Data show means ± SD. Each point in a-g represents data from individual mice. n values are the numbers of mice used to obtain the data. *P*-values were calculated with ordinary one-way ANOVA with Dunnett's post-hoc test (controls, NHA, HA-PHZ) or unpaired t-test (control-IgG and HA-TER119). The source data underlying Supplementary Fig. 1a-g, are provided as a Supplementary Data file.

**Supplementary Figure 2. Expansion of CECs in anemic mice**

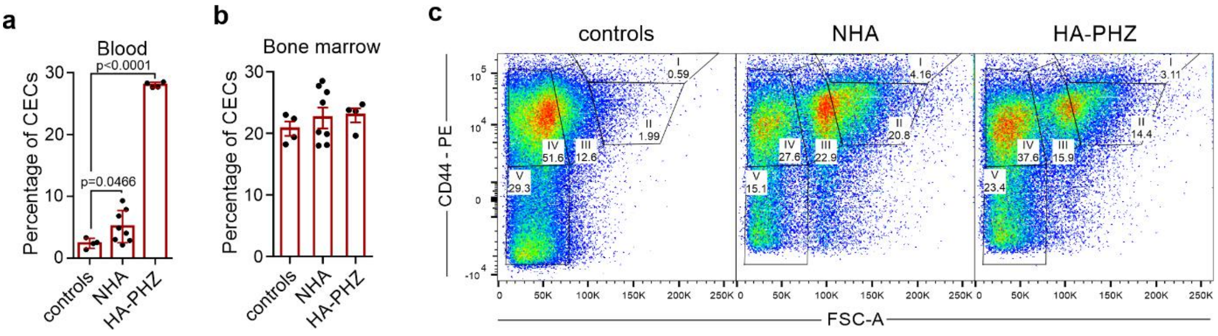


**a,** Percentages of CECs in the blood in anemic (NHA n=8, HA-PHZ n=4) and control mice (n=4). **b,** Percentages of CECs in the bone marrow of anemic (NHA n=8, HA-PHZ n=4) and control mice (n=4). *P*-value was calculated with Brown-Forsythe ANOVA test with Dunnett's T3 post-hoc test. **c,** Representative plot of CD44 and cell size (FSC) in CECs in healthy control mice and anemic (NHA, HA-PHZ) mice. Data show means ± SD. Each point in a-b represents data from individual mice. n values are the numbers of mice used to obtain the data. The source data underlying Supplementary Fig. 2a-b are provided as a Supplementary Data file.

**Supplementary Figure 3. Pro-inflammatory response by myeloid cells and induction of humoral immunity are not impaired in anemic mice**

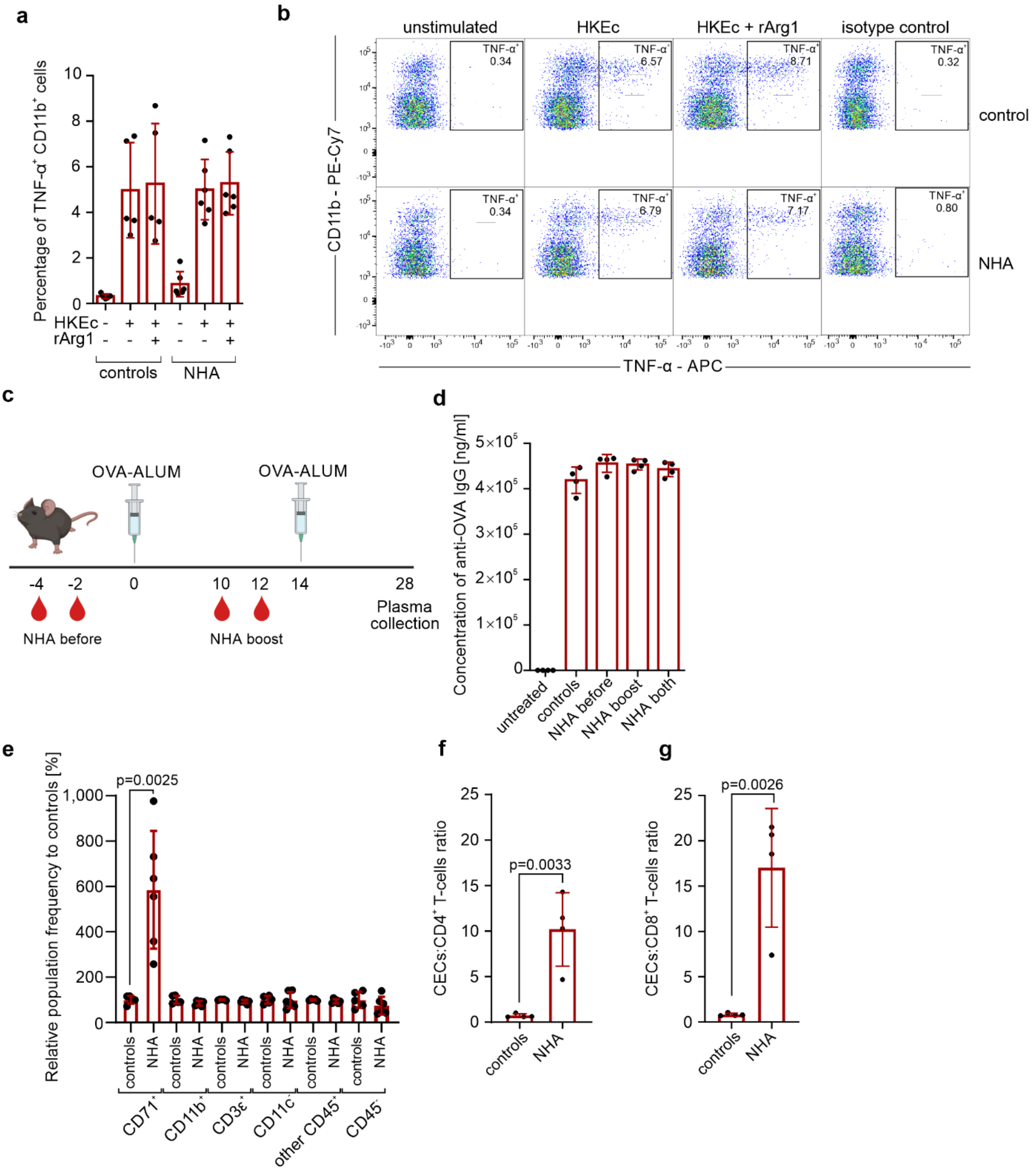


**a,** Splenocytes of control mice (n=5) and NHA mice (n=6) were stimulated with heat-killed *E. coli* (HKEc) for 6 hours in the presence of a protein transport inhibitor. TNF-α levels in CD11b$^+$ cells were determined by intracellular staining with fluorochrome-labeled antibodies. rARG1 – recombinant ARG1 (125 ng/ml). **b,** Representative plots of TNF-α levels in $CD11b^+$ cells from controls and NHA mice based on intracellular staining. **c,** Schematic presentation of the experimental setting. Non-hemolytic anemia (NHA) was induced in mice by phlebotomy before immunization (NHA before) with ovalbumin (OVA) with Alum adjuvant, before second immunization (NHA boost) or each time (NHA both). Blood was collected 28 days after first immunization and plasma was isolated for anti-OVA antibodies measurement. **d,** Concentrations of anti-OVA IgG antibodies in the plasma of controls (n=4) and NHA mice (each group n=4) were determined using ELISA. Untreated mice were not immunized with OVA (n=4). **e,** Relative population frequency in the spleen of anemic mice (NHA, n=6) compared to healthy controls (n=5). **f,** Ratio of CECs to $CD4^+$ T-cells in the spleen of control (n=5) and anemic mice (n=5). **g** Ratio of CECs to $CD8^+$ T-cells in the spleen of control (n=5) and anemic mice (n=5). *P*-value was calculated with an unpaired *t*-test. Data show means ± SEM (a) or means ± SD (d-g). Each point in a, d, e, f, g represents data from individual mice. n values are the numbers of mice used to obtain the data. The source data underlying Supplementary Fig. 3a, Supplementary Figure 3d-g are provided as a Supplementary Data file.

**Supplementary Figure 4. The purity of isolated murine CECs**

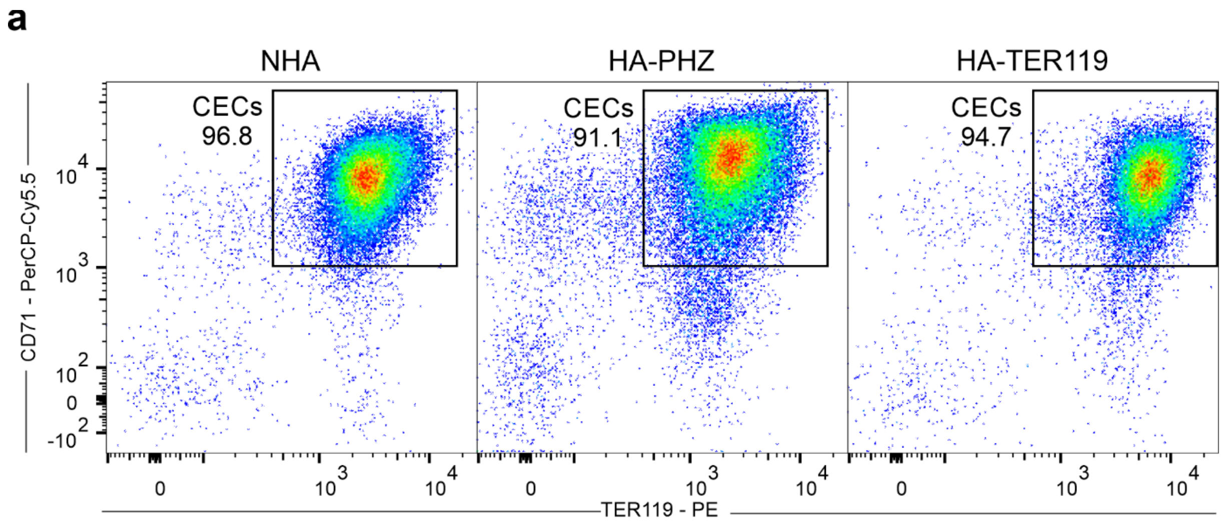


**a,** Representative plots of CD71 and TER119 in CECs isolated from the spleens of anemic mice (NHA, HA-PHZ, HA-TER119) using positive immunomagnetic selection with anti-CD71 antibody.

**Supplementary Figure 5. The level of reactive oxygen species (ROS) is the highest in early-stage CECs**

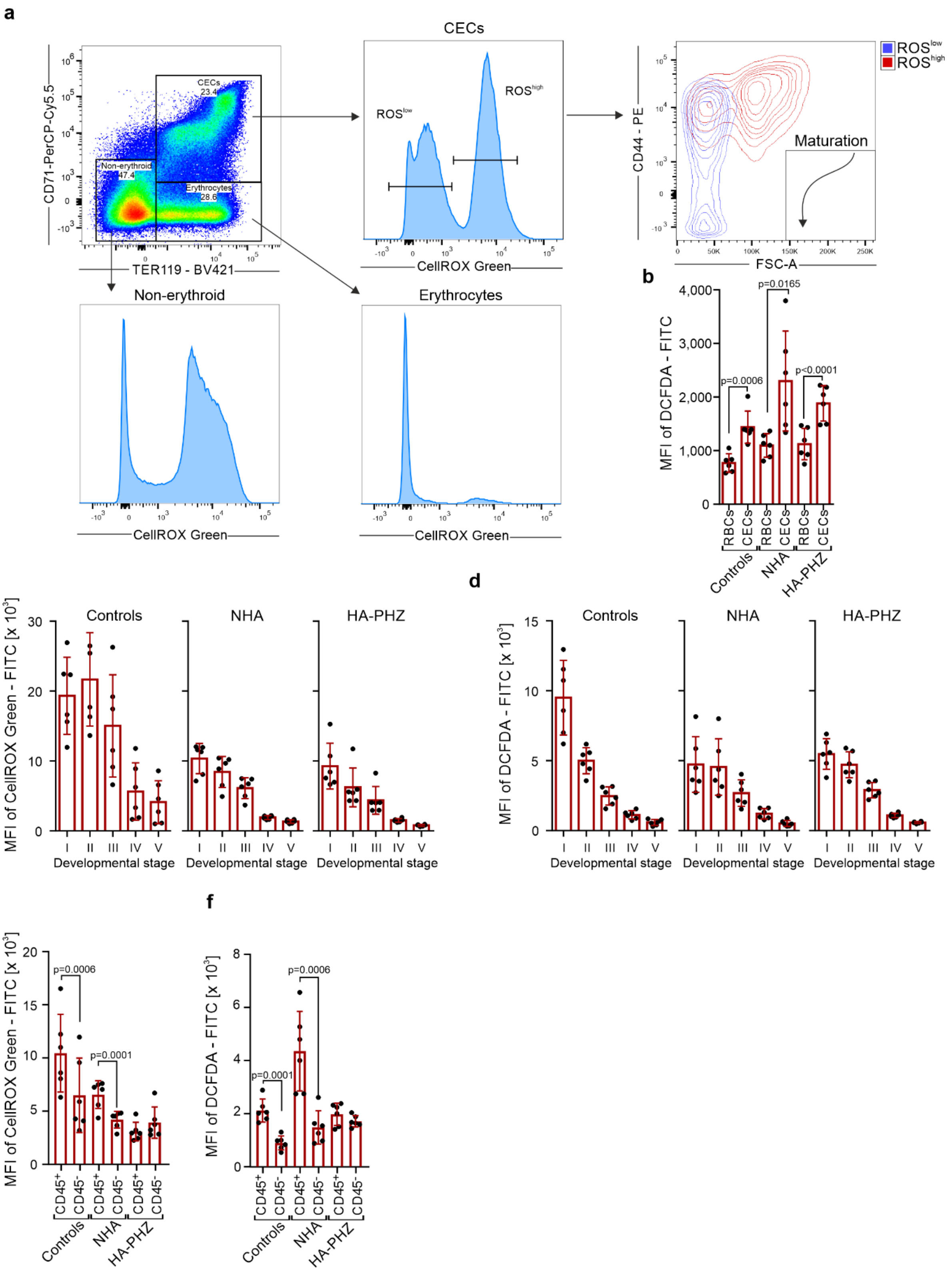

**a,** Representative plot of CD71 and TER119 levels in live cells and histograms of reactive oxygen species (ROS) levels based on CellROX Green – FITC in CECs ($CD71^{+}TER119^{+}$), erythrocytes (RBCs, $CD71^{-}TER119^{+}$), and non-erythroid cells ($CD71^{-}TER119^{-}$). **b,** ROS levels in CECs and RBCs in the spleen based on mean fluorescence intensity (MFI) 2',7'-dichlorodihydrofluorescein diacetate (DCFDA) – FITC in anemic (NHA n=6, HA-PHZ n=6) and control mice (n=6). *P*-values were calculated with an unpaired *t*-test. **c,d,** ROS levels in CECs at different stages of differentiation based on mean fluorescence intensity (MFI) of CellRox Green – FITC (**c**) and 2',7'-dichlorodihydrofluorescein diacetate (DCFDA) – FITC (**d**) in anemic (NHA n=6, HA-PHZ n=6) and control mice (n=6). **e,f,** ROS levels in $CD45^{+}$ CECs and $CD45^{-}$ CECs based on mean fluorescence intensity (MFI) of CellRox Green – FITC (**e**) and 2',7'-dichlorodihydrofluorescein diacetate (DCFDA) – FITC (**f**) in anemic (NHA n=6, HA-PHZ n=6) and control mice (n=6). *P* values were calculated with paired *t*-test. Data show means ± SD. Each point in b-f represents data from individual mice. n values are the numbers of mice used to obtain the data. The source data underlying Supplementary Fig. 5b-f are provided as a Supplementary Data file.

**Supplementary Figure 6. ARG1 and ARG2 levels are the highest in the early-stages CECs**

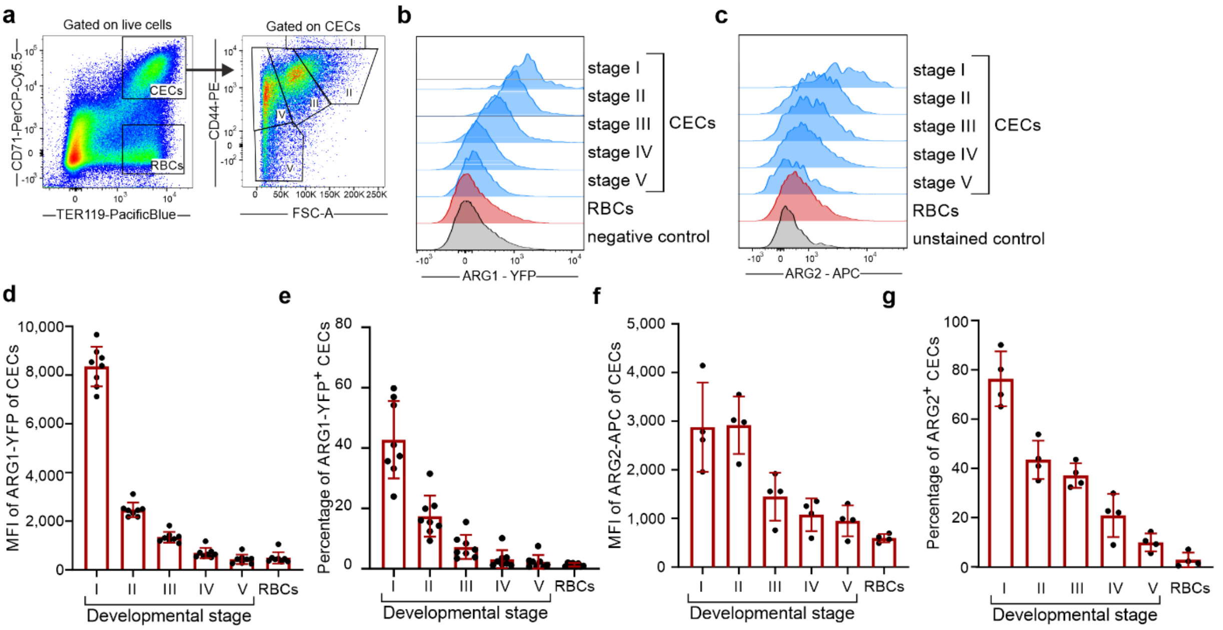


**a,** Representative plot of CD71 and TER119 levels in live cells in the spleens of NHA mice and gating strategy of CECs developmental stages based on CD44 levels and relative cell size. **b,** Representative histograms of ARG1 expression in different developmental stages of CECs and RBCs in NHA mouse based on YFP mean fluorescence intensity (MFI) in reporter B6.129S4-Arg1$^{tm1Lky}$/J mice. Negative control represents the background fluorescence of YFP in wild-type C57Bl/6 mouse. **c,** Representative histograms of ARG2 expression in different developmental stages of CECs and RBCs based on intracellular staining in NHA mouse. Unstained control represents sample unstained for ARG2. **d,e,** ARG1 expression (**d**) and percentage of ARG1$^{+}$ cells (**e**) in different developmental stages of CECs in NHA mice based on YFP mean fluorescence intensity (MFI) in reporter B6.129S4-Arg1$^{tm1Lky}$/J mice (n=8). **f,g,** ARG2 levels (**f**) and percentage of ARG2$^{+}$ cells (**g**) in different developmental stages of CECs in NHA mice based on intracellular staining (n=4). Data show means ± SD. Each point in d-g represents data from individual mice. n values are the numbers of mice used to obtain the data. The source data underlying Supplementary Fig. 6d-g are provided as a Supplementary Data file.

**Supplementary Figure 7. ARG1 and ARG2 expression in CECs of anemic mice**

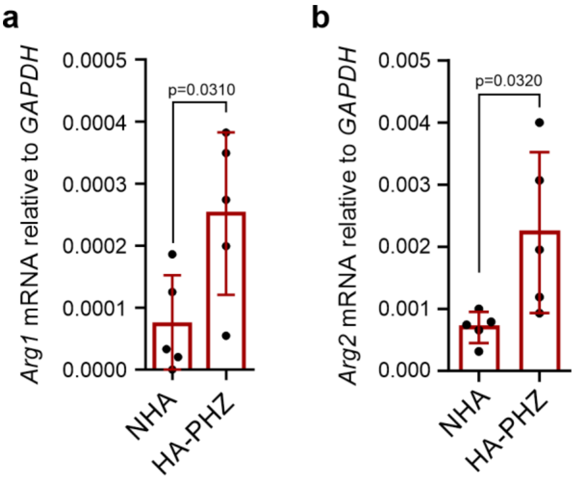


**a,b,** mRNA expression of ARG1 (**a**) and ARG2 (**b**) in CECs isolated from NHA (n=5) and HA-PHZ (n=5) mice was determined using real-time quantitative PCR method. *P*-values were calculated with an unpaired *t*-test. Data show means ± SD. Each point in a, b represents data from individual mice. n values are the numbers of mice used to obtain the data. The source data underlying Supplementary Fig. 7a,b are provided as a Supplementary Data file.

**Supplementary Figure 8. PHZ diminishes immunoregulatory functions of murine CECs *ex vivo***

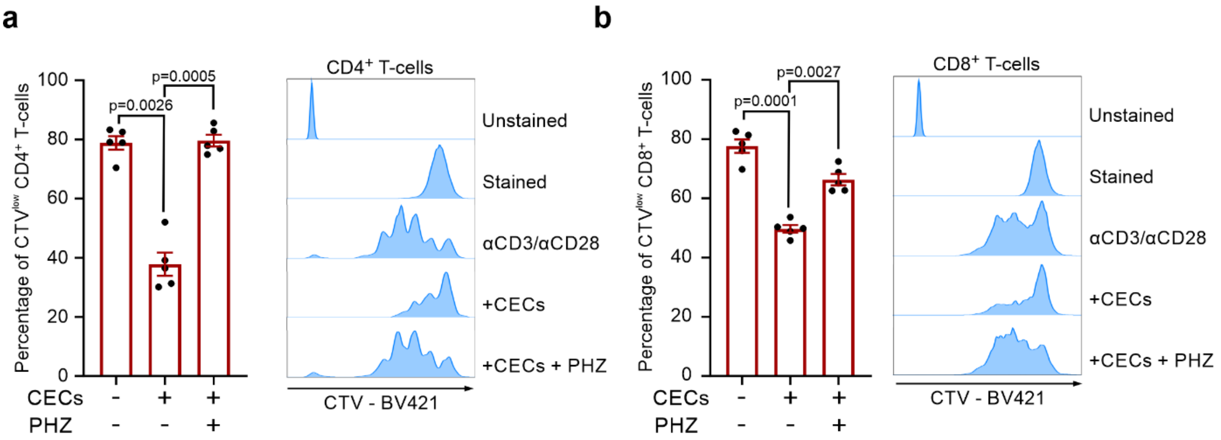


**a,b,** Effects of phenylhydrazine (PHZ, 100 µM) on the proliferation of CD4$^{+}$ (**a**) and CD8$^{+}$ (**b**) T-cells stimulated with αCD3/αCD28 and co-cultured with CECs at a 1:2 ratio (T-cells:CECs) isolated from the spleens of NHA mice (n=5). Representative proliferation histograms of αCD3/αCD28-stimulated CD4$^{+}$ (**a**) and CD8$^{+}$ (**b**) T-cells co-cultured with CECs in the presence of PHZ. Histograms show the fluorescence of CTV (CellTraceViolet) – BV421. *P*-values were calculated with repeated-measures ANOVA with Tukey's post-hoc test. Data show means ± SEM. Each point in a represents data from individual mice. n values are the numbers of mice used to obtain the data. The source data underlying Supplementary Fig. 8a,b are provided as a Supplementary Data file

**Supplementary Figure 9. Molecular analysis of arginases – PHZ complexes**

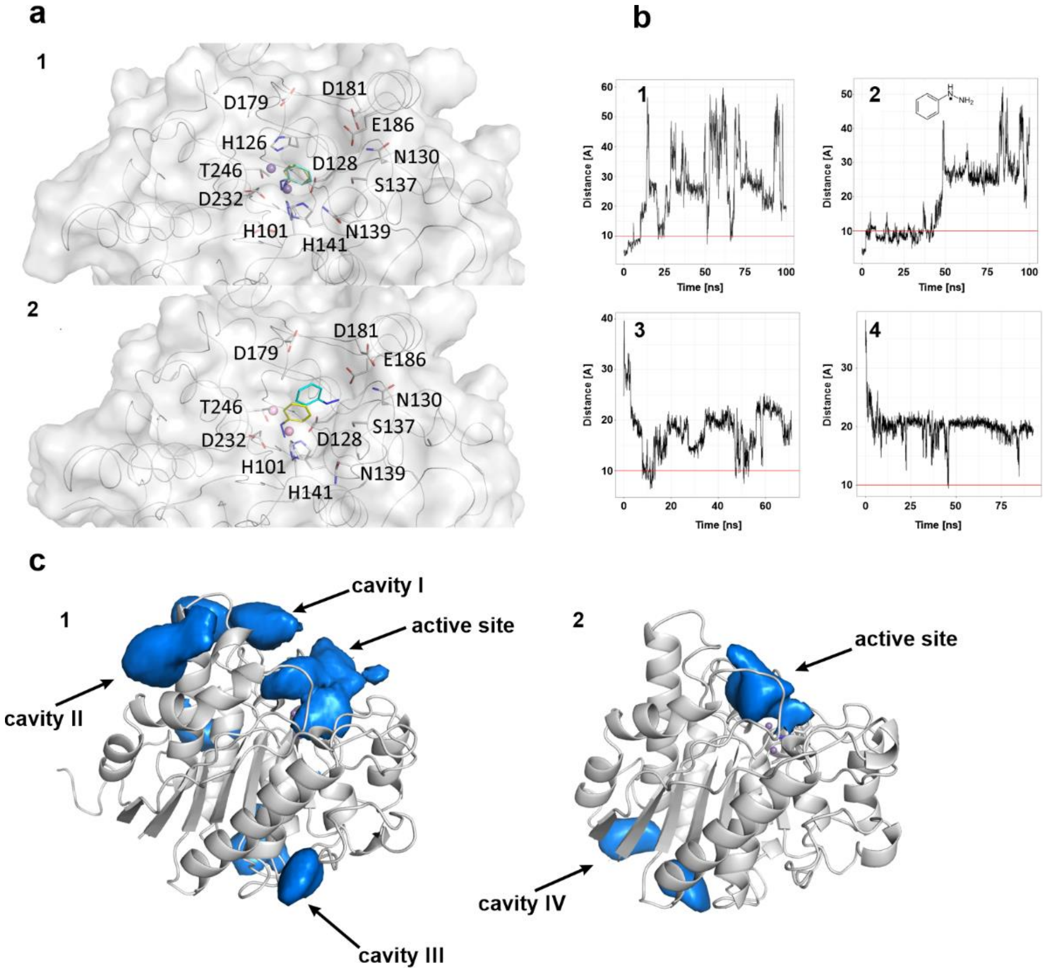


**a**, The predicted conformation of PHZ in complex with human (panel 1) and mouse (panel 2) ARG1. The best-scored conformations of the ligand proposed independently by surflex and GOLD are colored cyan and yellow, respectively. Sidechains of key active site residues are also depicted. **b,** The distance between carboxyl group of D126 and benzene-substituted nitrogen atom of PHZ was observed during 100 ns MD simulations. (1) PHZ and human ARG1 (2) PHZ and human ARG2. In both cases the ligand was initially present in the active site. (3) The same distance but for MD simulations involving human ARG1 and six molecules of PHZ randomly placed around the enzyme. The distance for a molecule closest to the active site at a given moment in time is reported. (4) Same as 3 but for ARG2. The vertical red line marks 10 Å value. If the distance is below that threshold we assume the ligand can effectively shield the active site from other molecules **c,** During MD simulation involving PHZ initially placed in the active site of ARG1 (1) and ARG2 (2) the ligand binds to several other regions on the surface of the protein.

**Supplementary Figure 10. Reactive oxygen species scavengers do not prevent ARG1 induction by PHZ.**

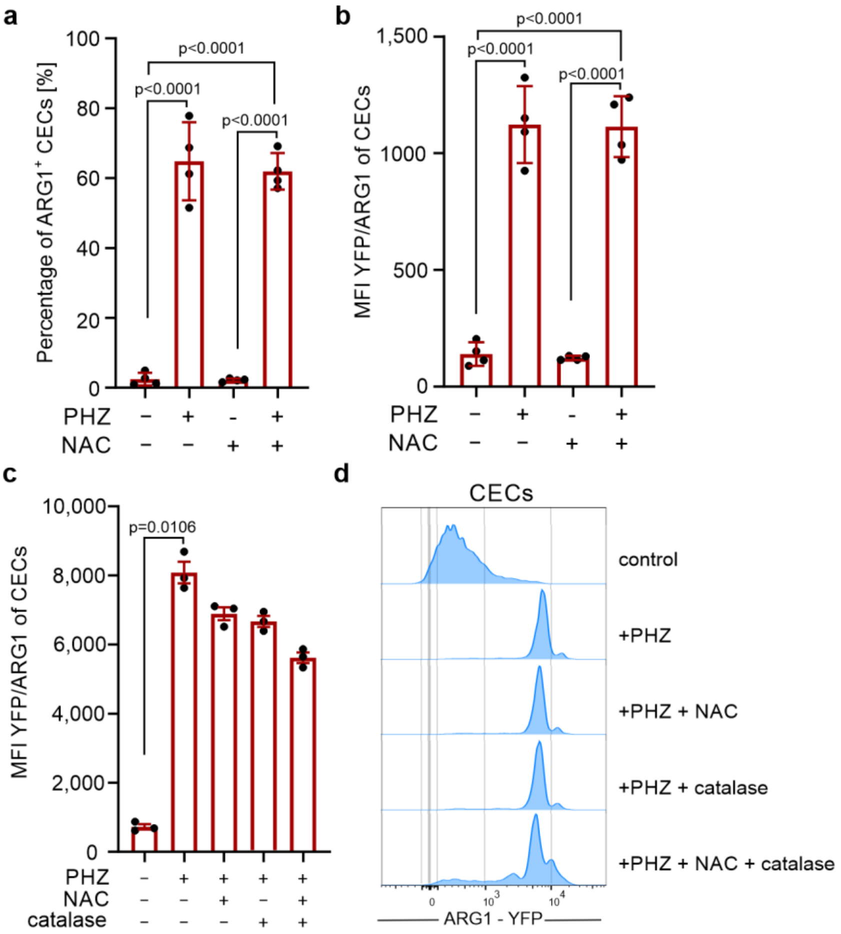


**a-b**, YARG mice were injected intraperitoneally (i.p.) three days before harvest with 50 mg per kg body weight of phenylhydrazine (PHZ) and three times every 24h with 150 mg per kg body weight of N-acetylcysteine (NAC) (each group n=4). Percentages of ARG1$^+$ CECs (**a**) and mean fluorescence intensity (MFI) of YFP-ARG1 (**b**). **c,d**, CECs were isolated from the spleens of NHA YARG mice and incubated with 100 µM phenylhydrazine (PHZ), 100 µM NAC, and/or 100 µg/ml catalase. The fluorescence of YFP/ARG1 was assessed after 24h by flow cytometry (**c**). Histograms show representative fluorescence of ARG1 – YFP of CEC (**d**). *P*-values were calculated with repeated-measures ANOVA with Bonferroni's post-hoc test. Data show means ± SD (a-b) or means ± SEM (c). Each point in a-c represents data from individual mice. n values are the numbers of mice used to obtain the data. The source data underlying Supplementary Fig. 10a-c are provided as a Supplementary Data file.

**Supplementary Figure 11. Neonatal CECs suppress T-cells proliferation by arginase and ROS**

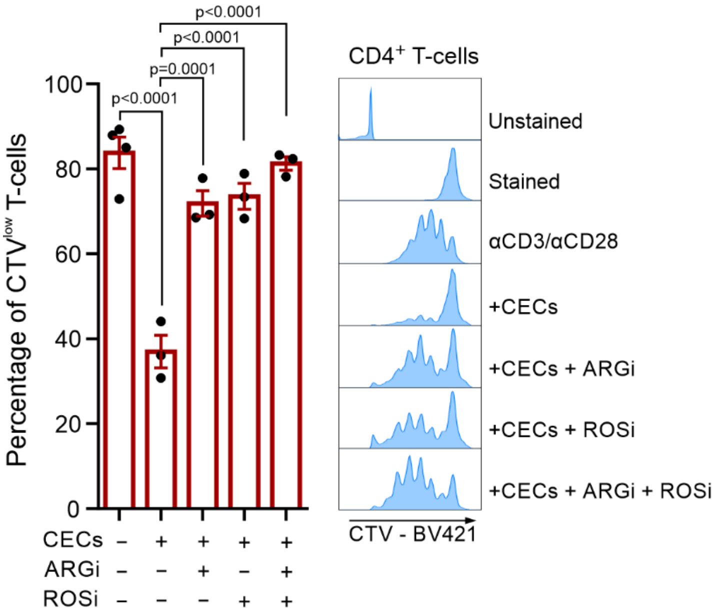


Effects of ARGi (OAT-1746, 500 nM) and ROSi (N-acetylcysteine, 100 μM) on the proliferation of αCD3/αCD28-stimulated $CD4^+$ T-cells co-cultured with CECs in a ratio of 1:2 isolated from the spleens of neonatal mice (n=3). Representative proliferation histograms of αCD3/αCD28-stimulated $CD4^+$ T-cells co-cultured with CECs in the presence of ARGi or ROSi. Histograms show the fluorescence of CTV (CellTraceViolet) – BV421. *P*-values were calculated with one-way ANOVA with Bonferroni's post-hoc test. Data show means ± SEM. Each point represents data from individual mice. n values are the numbers of mice used to obtain the data. The source data underlying Supplementary Fig. 11 are provided as a Supplementary Data file

**Supplementary Figure 12. Anemic mice have slightly decreased L-arginine concentration in serum**

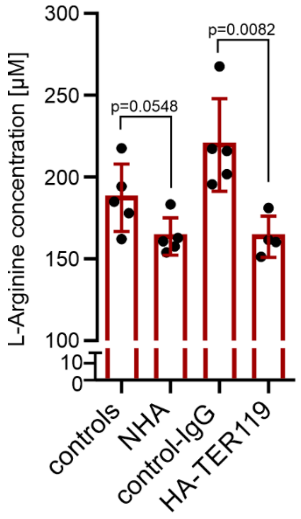


L-Arginine concentration in the serum of anemic and healthy mice determined by mass spectrometry (controls n=5, NHA n=5, control-IgG n=5, HA-TER119 n=4). *P*-values were calculated using unpaired *t*-tests. Data show means ± SD. Each point in a represents data from individual mice. n values are the numbers of mice used to obtain the data. The source data underlying Supplementary Fig. 12 are provided as a Supplementary Data file.

**Supplementary Figure 13. Lymph node T-cells have normal CD3ζ levels in anemic mice**

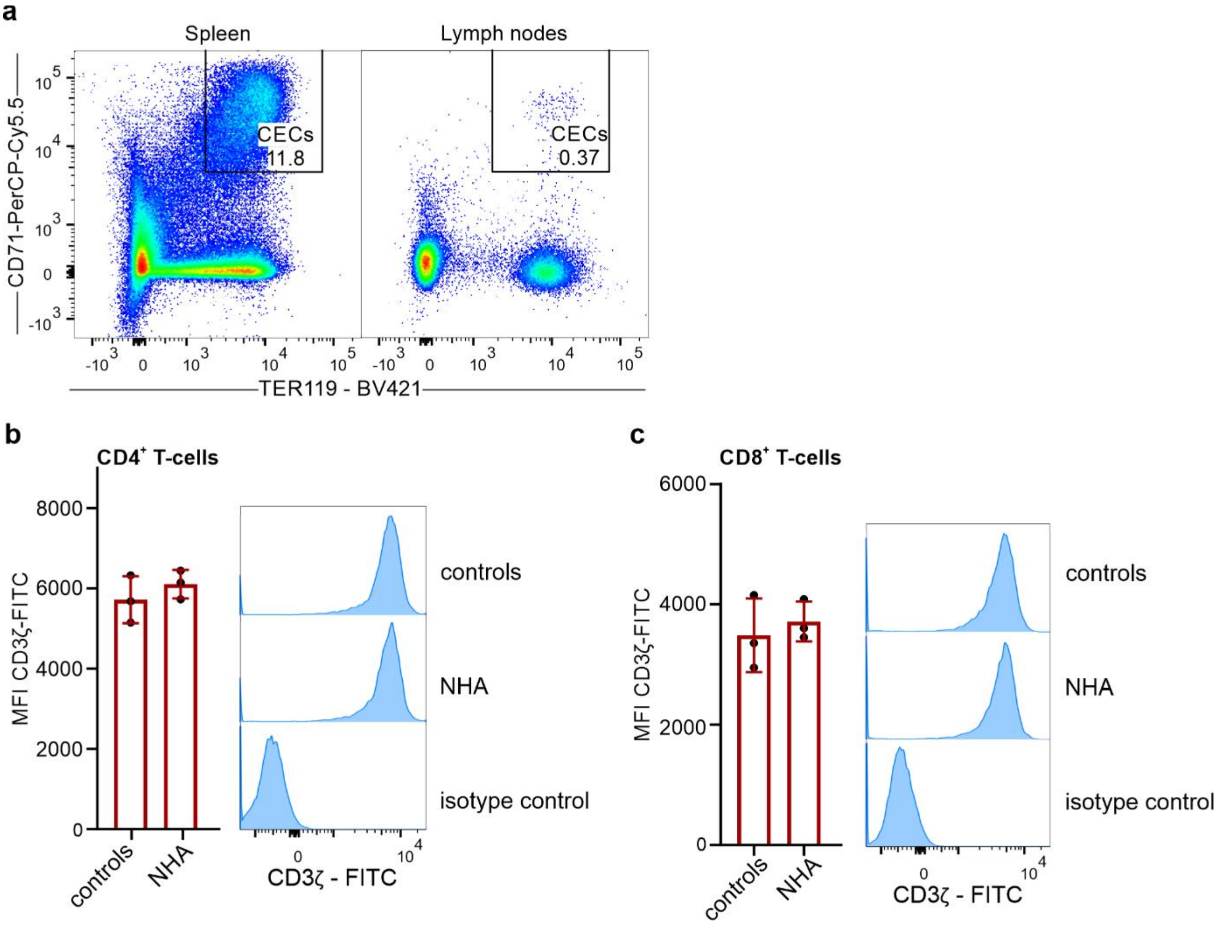


**a,** Representative plots of CD71 and TER119 in live cells from the spleen as well as inguinal and axillary lymph nodes isolated from NHA mice. **b,c,** The levels of CD3ζ in the lymph node CD4$^{+}$ (**b**) and CD8$^{+}$ T-cells (**c**) of controls (n=3) and anemic (n=3) mice. Histograms show the fluorescence of CD3ζ – FITC. *P*-values were >0.05. *P*-value was calculated with the Mann-Whitney test. Data show means ± SD. Each point in b, c represents data from individual mice. n values are the numbers of mice used to obtain the data. The source data underlying Supplementary Fig. 13b,c are provided as a Source Data file.

### Supplementary Figure 14. Stress erythropoiesis and Arg1 expression in *Arg2*$^{-/-}$ mice

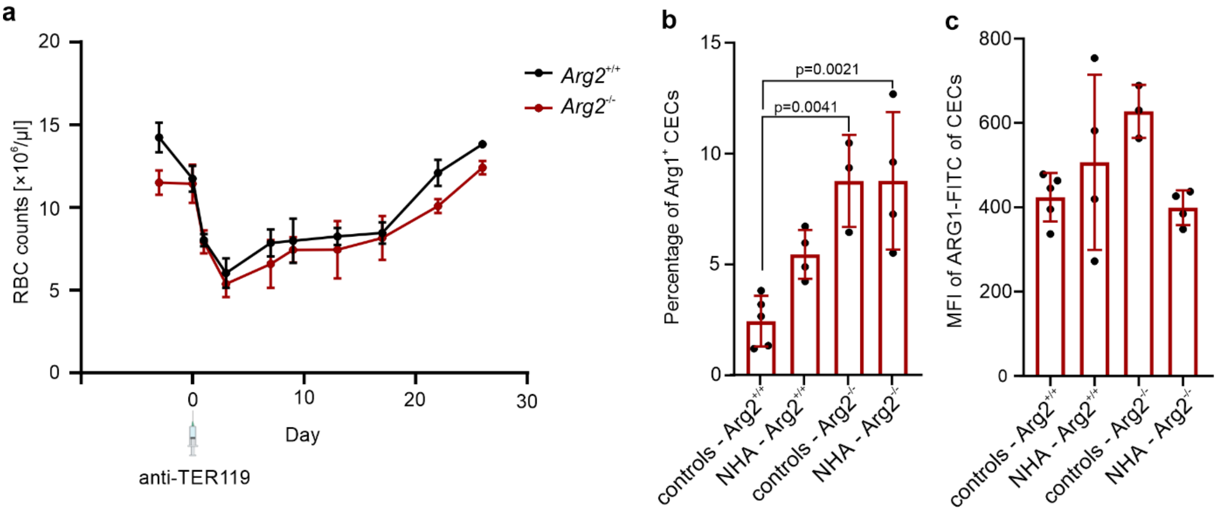


**a,** *Arg2*$^{-/-}$ knockout mice (n=3) and wild-type mice (n=3) were administered with 30 µg anti-TER119 antibody to induce stress erythropoiesis. Red blood cell (RBC) count was determined using an automated hematology analyzer. $P > 0.05$ for the analysis of the differences between *Arg2*$^{-/-}$ knockout mice and wild-type *Arg2*$^{+/+}$ mice in all time points. *P* values were calculated with the Mann-Whitney test. **b,** Percentages of ARG1$^{+}$ CECs in the spleens of control (n=5), anemic wild-type (n=4), *Arg2*$^{-/-}$ knockout anemic (n=4), and control mice (n=3). **c,** ARG1 expressison in CECs based on intracellular staining in the spleen of control (n=5), anemic wild-type (n=4), *Arg2*$^{-/-}$ knockout anemic (n=4), and control mice (n=3). *P*-values were calculated with one-way ANOVA with Bonferroni's post-hoc test. Data show means ± SD. Each point in b, c represents data from individual mice. n values are the numbers of mice used to obtain the data. The source data underlying Supplementary Fig. 14a-c are provided as a Source Data file.

**Supplementary Figure 15. T-cells proliferation and TNF-α production by myeloid cells is unimpaired in PBMCs of anemic patients**

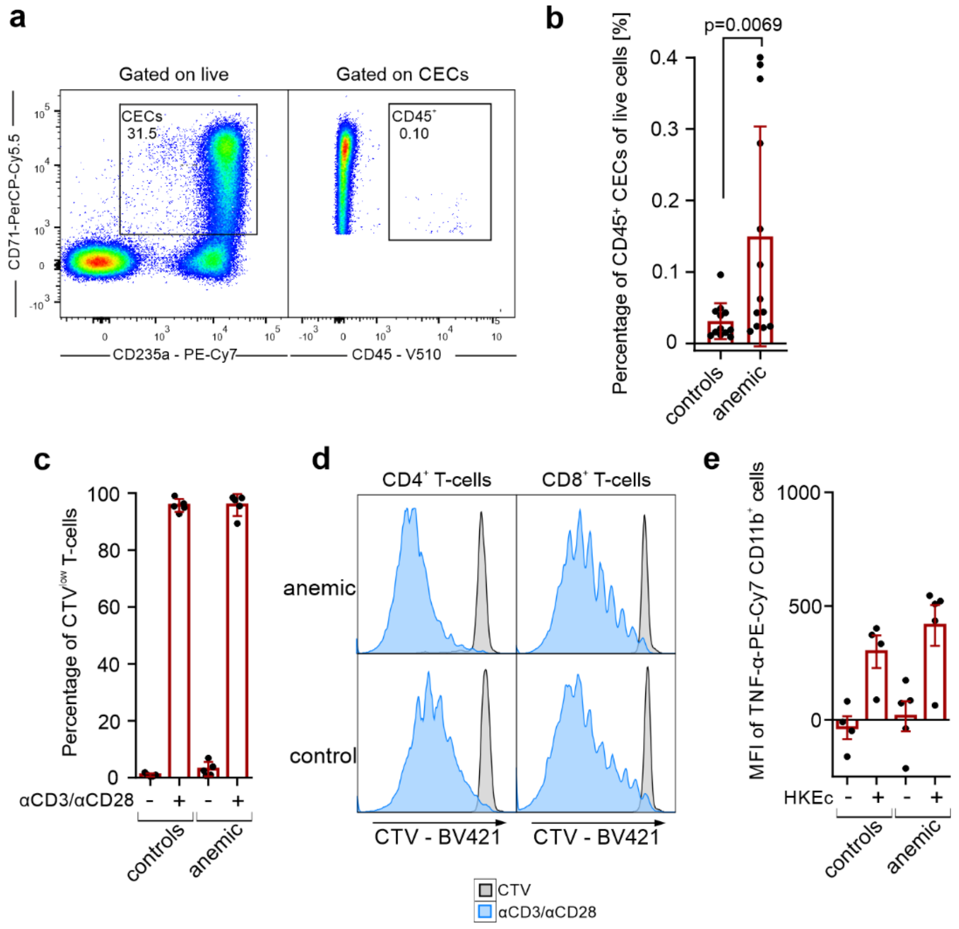


**a,** Representative plots of CD71 and CD235a levels in PBMCs and CD45 levels in CECs from an anemic individual. **b,** Percentages of live CD45$^+$ CECs cells in PBMCs of controls (n=12) and anemic patients (n=13). **c,** Proliferation of αCD3/αCD28-stimulated CD3ε$^+$ T-cells in PBMCs of controls (n=5) and anemic patients (n=5). **c,** Representative proliferation histograms of αCD3/αCD28-stimulated PBMC from anemic or control patients. **d,** Representative histograms of the fluorescence of CTV (CellTraceViolet) – BV421 in αCD3/αCD28-stimulated CD4$^+$ and CD8$^+$ T-cells. **e**, PBMCs of control patients (n=4) and anemic patients (n=5) were stimulated with Heat-killed *E. coli* (HKEc) for 12 hours in the presence of a protein transport inhibitor. TNF-α levels in CD11b$^+$ cells were determined by intracellular staining with fluorochrome-labeled antibodies. *P* values were calculated using the Mann-Whitney test (**b,c,d**). Data show means ± SD (**b**) or means ± SEM (**c,e**). Each point in **b, c, e** represents data from individual patients. n values are the numbers of individual patients used to obtain the data. The source data underlying Supplementary Fig. 15b,c,e are provided as a Source Data file.

**Supplementary Figure 16. CECs in the human bone marrow are enriched in early-stage progenitors and express ARG1 and ARG2**

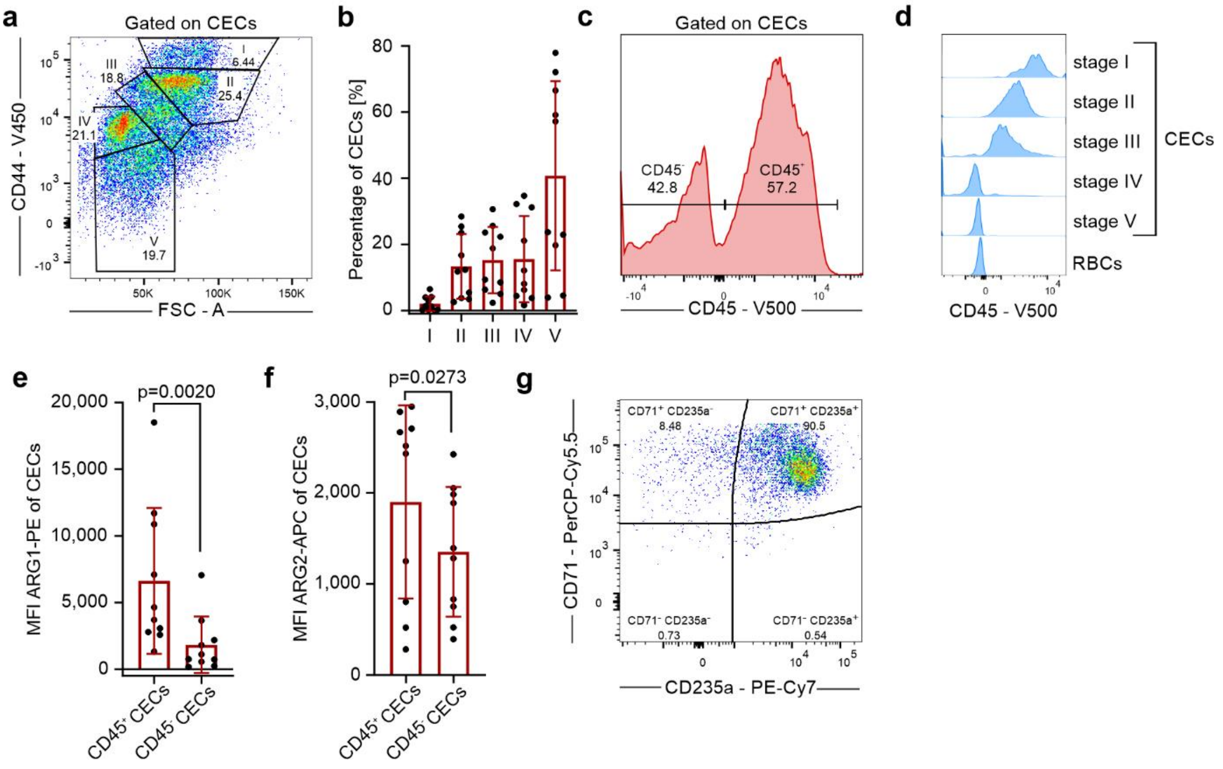


**a,** Representative dot plot of the gating of CECs developmental stages based on CD44 levels and relative cell size (FSC-A) in CECs from human bone marrow. **b,** Percentages of CECs in different developmental stages in human bone marrow (n=10). **c,** Representative histogram of CD45 levels in CECs from human bone marrow. **d,** Representative histogram of CD45 levels in CECs at different developmental stages. Red blood cells (RBCs) are shown as CD45-negative control. **e,f,** The levels of ARG1 (**e**) and ARG2 (**f**) in $CD45^+$ and $CD45^-$ CECs from human bone marrow. **g,** Representative plot of CD71 and CD235a in isolated CECs from human bone marrow. *P* values were calculated using the Mann-Whitney test (**d,e**). Data show means ± SD. Each point in **b, e, f** represents data from individual patients. n values are the numbers of individual patients used to obtain the data. The source data underlying Supplementary Fig. 16b,e,f are provided as a Source Data file.

**Supplementary Figure 17. Erythroleukemia-derived erythroid cell lines have high arginase activity and ROS levels**

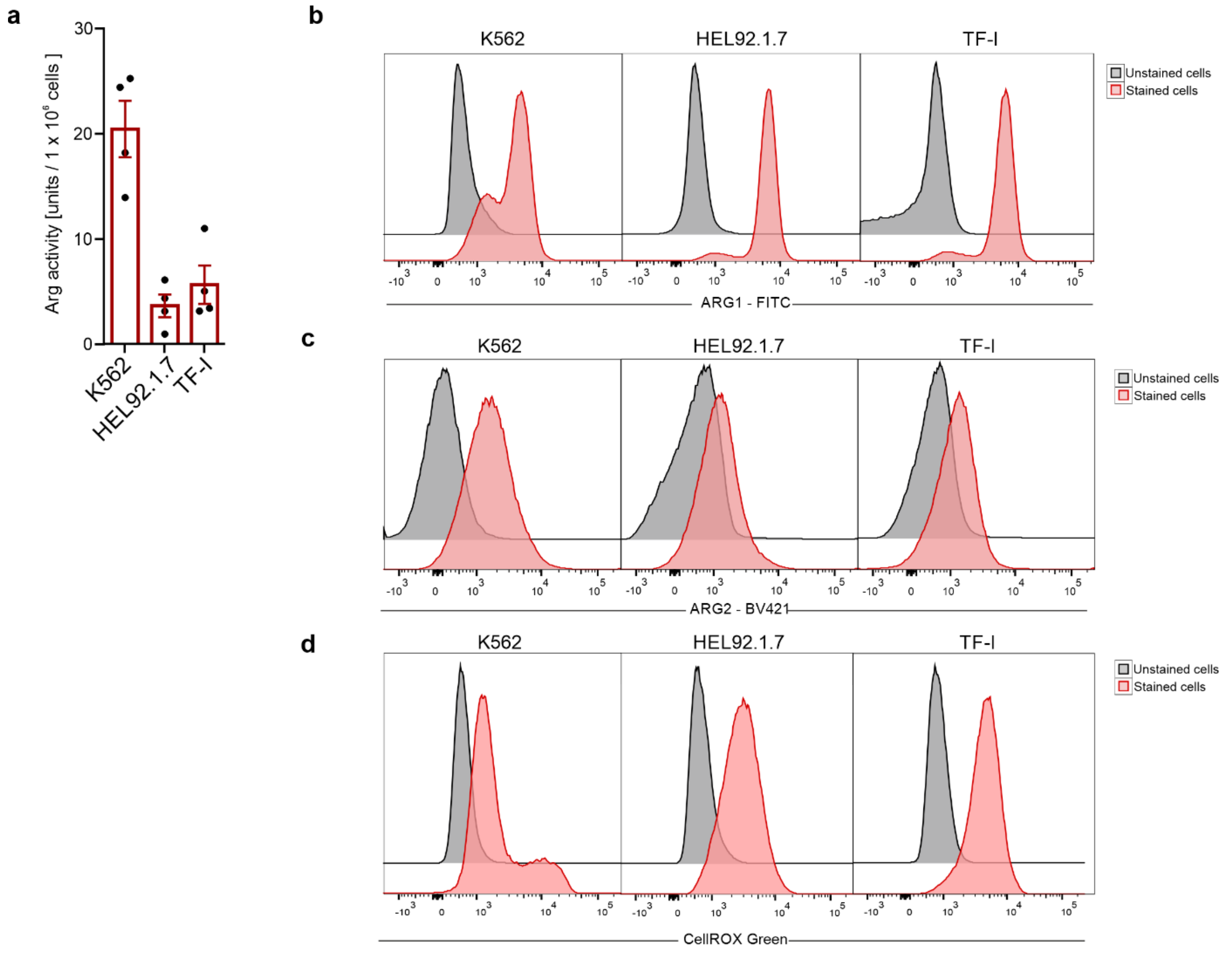


**a,** Arginase activity in erythroid cells calculated per $1{\times}10^{6}$ cells (n=4). **b,c,** Representative histograms of ARG1 (**b**) and ARG2 (**c**) expression in erythroid cell lines. **e,** Representative histograms of ROS levels based on CellRox Green – FITC fluorescence in erythroid cell lines. Data show means ± SEM. Each point in **a** represents data from one biological replicate. n values are the numbers of biological replicates used to obtain the data. The source data underlying Supplementary Fig. 17a are provided as a Source Data file.

**Supplementary Figure 18. Erythroid cells suppress T-cells in an ARG- and ROS-dependent mechanism**

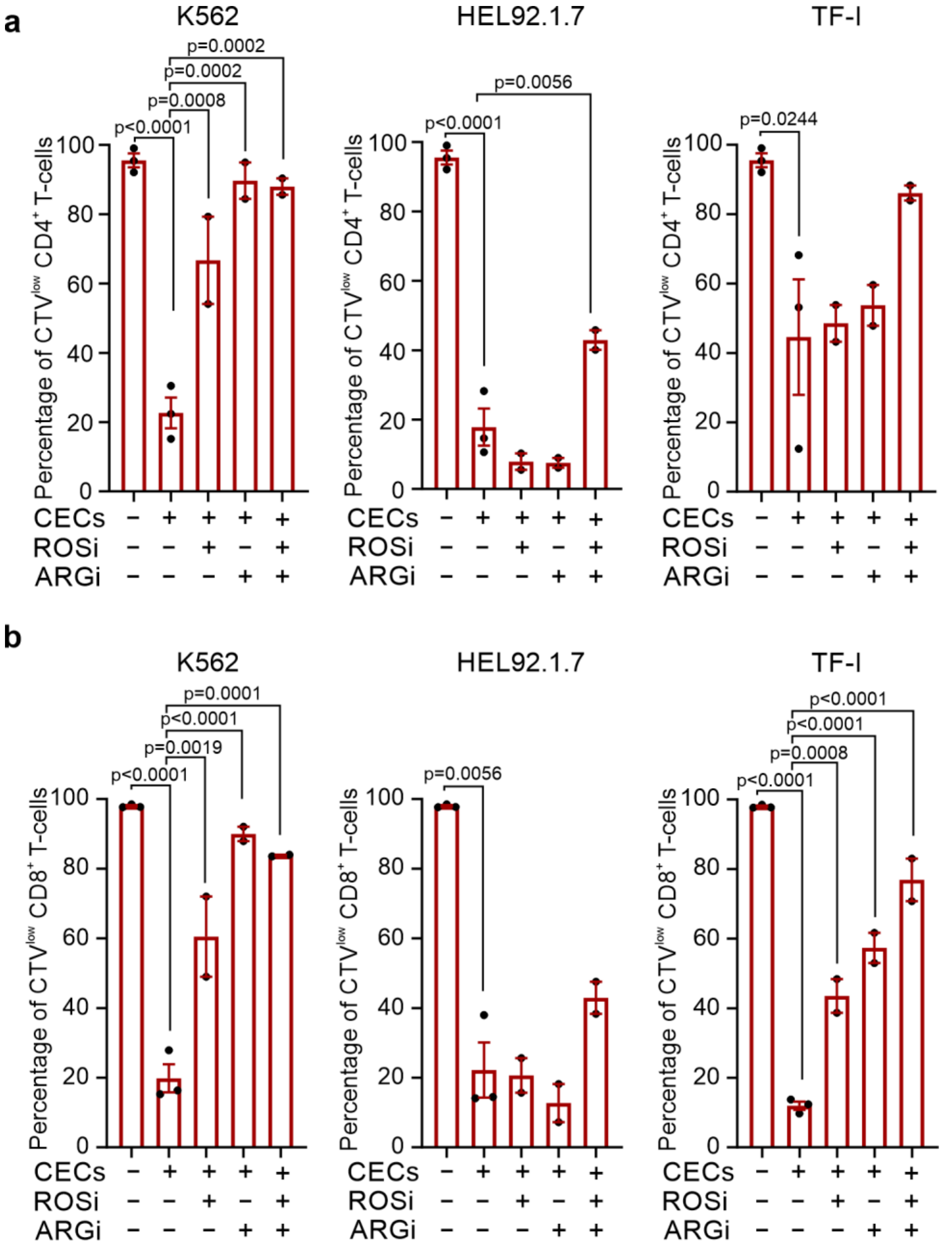


**a,b,** Effects of ARGi (OAT-1746, 1.5 μM) and ROSi (N-acetylcysteine, 200 μM) on proliferation of CTV-labelled $CD4^+$ (**a,** n=2) and $CD8^+$ (**b,** n=2) T-cells triggered by αCD3/αCD28 and co-cultured with erythroid cell lines at a 1:2 ratio. Data correspond to Fig. 9e (**a**) and 9i (**b**). Representative histograms are shown in Fig. 9e and Fig. 9i. *P* values were calculated with ordinary one-way ANOVA with Holm-Sidak's post-hoc test. Data show means ± SEM. Each point in a, b represents data from a mean of technical duplicates. n values are the numbers of biological repetitions of *in vitro* experiments. The source data underlying Supplementary Fig. 18a,b are provided as a Source Data file.

**Supplementary Figure 19. CECs expansion and differentiation from PBMC**

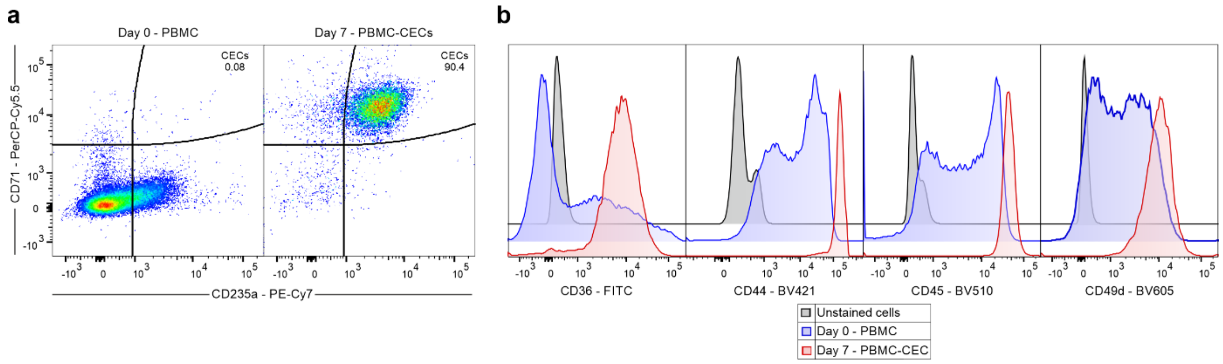


**a,** Representative plots of CD71 and CD235a expression in PBMCs on days 0 and 7 after erythroid differentiation. **b,** Representative histograms of CD36, CD44, CD45, and CD49d levels in PBMC and CECs differentiated from PBMCs (PBMC-CECs).

**Supplementary Figure 20. PBMC-CECs had high ARG1 and ARG2 expression**

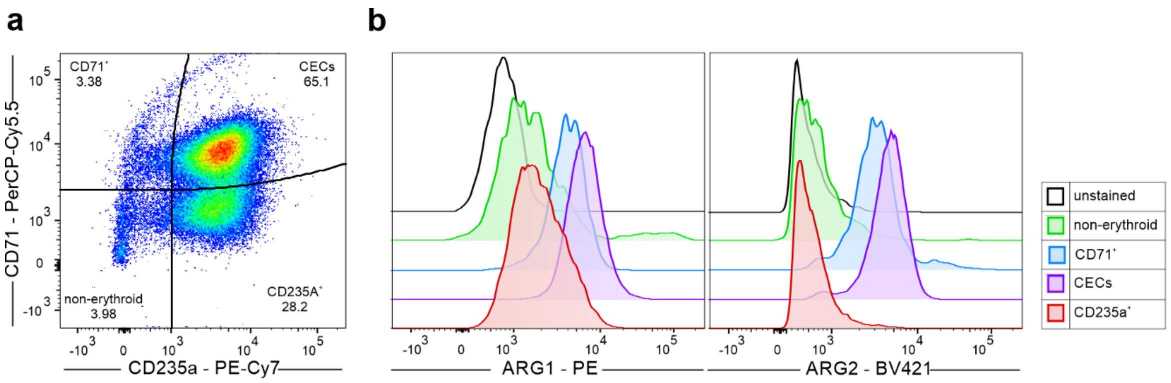


**a,** Representative plots of CD71 and CD235a expression in CECs differentiated from PBMC. **b,** Histograms of ARG1 and ARG2 expression in CECs ($CD71^{+}CD235a^{+}$), $CD71^{+}$ cells, RBCs ($CD71^{-}CD235a^{+}$) and non-erythroid cells ($CD71^{-}CD235a^{-}$).

**Supplementary Figure 21. PBSCs have high ARG1 and ARG2 expression but do not suppress T-cells proliferation**

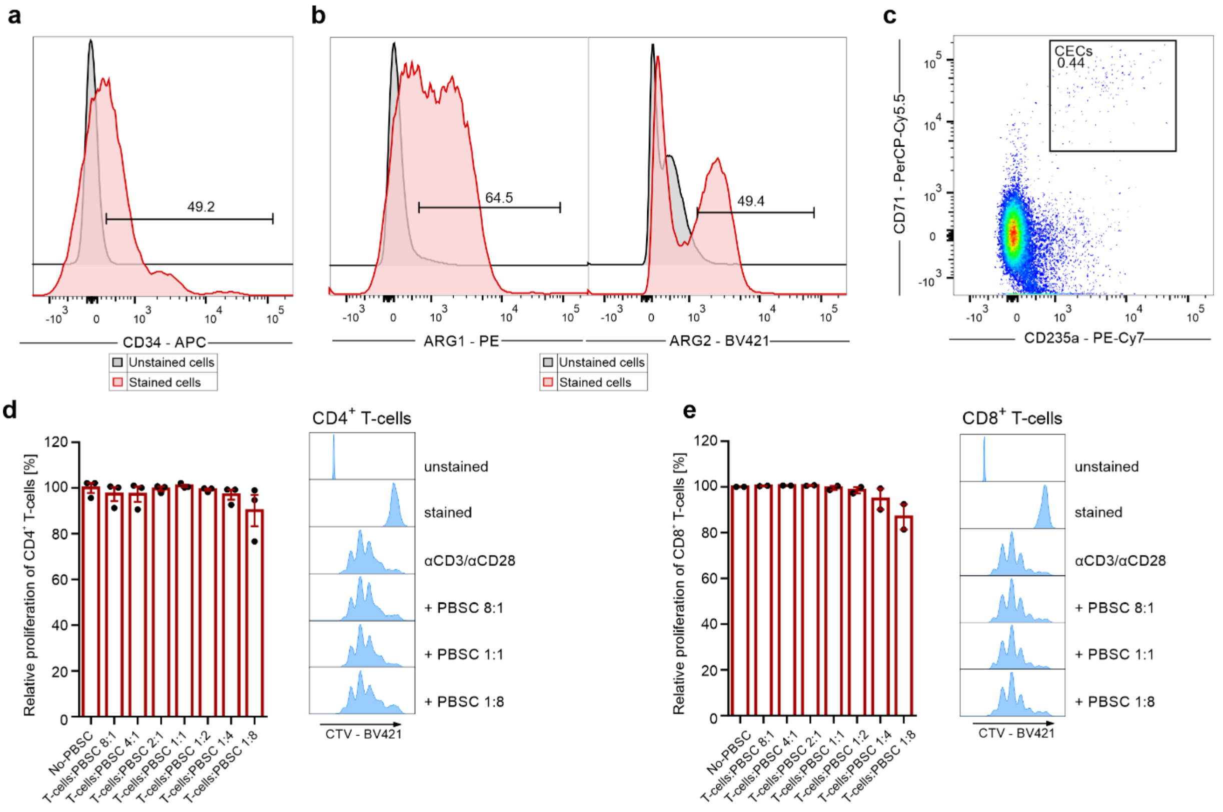


**a,** Representative histogram showing CD34 levels in peripheral blood stem cells (PBSC). **b,** Representative histograms of ARG1 and ARG2 expression in PBSCs. **c,** Representative plot of CD71 and CD235a expression in PBSCs. **d,e,** Proliferation of CTV-labelled CD4$^+$ (**d,** n=3) and CD8$^+$ (**e,** n=2) T-cells triggered by αCD3/αCD28 and co-cultured with PBSC. Relative proliferation was calculated with No-PBSC T-cells as a control. Representative histograms show the fluorescence of CTV (CellTraceViolet) – BV421 in CD4$^+$ (**d**) and CD8$^+$ T-cells (**e**) stimulated with αCD3/αCD28. Data show means ± SEM. Each point in **d**, **e** represents data from an individual patient. n values are the numbers of patients used to obtain the data. The source data underlying Supplementary Fig. 21d,e are provided as a Source Data file.

### Supplementary Figure 22. PBMC-derived CECs suppress T-cells in ARG- and ROS-dependent mechanism

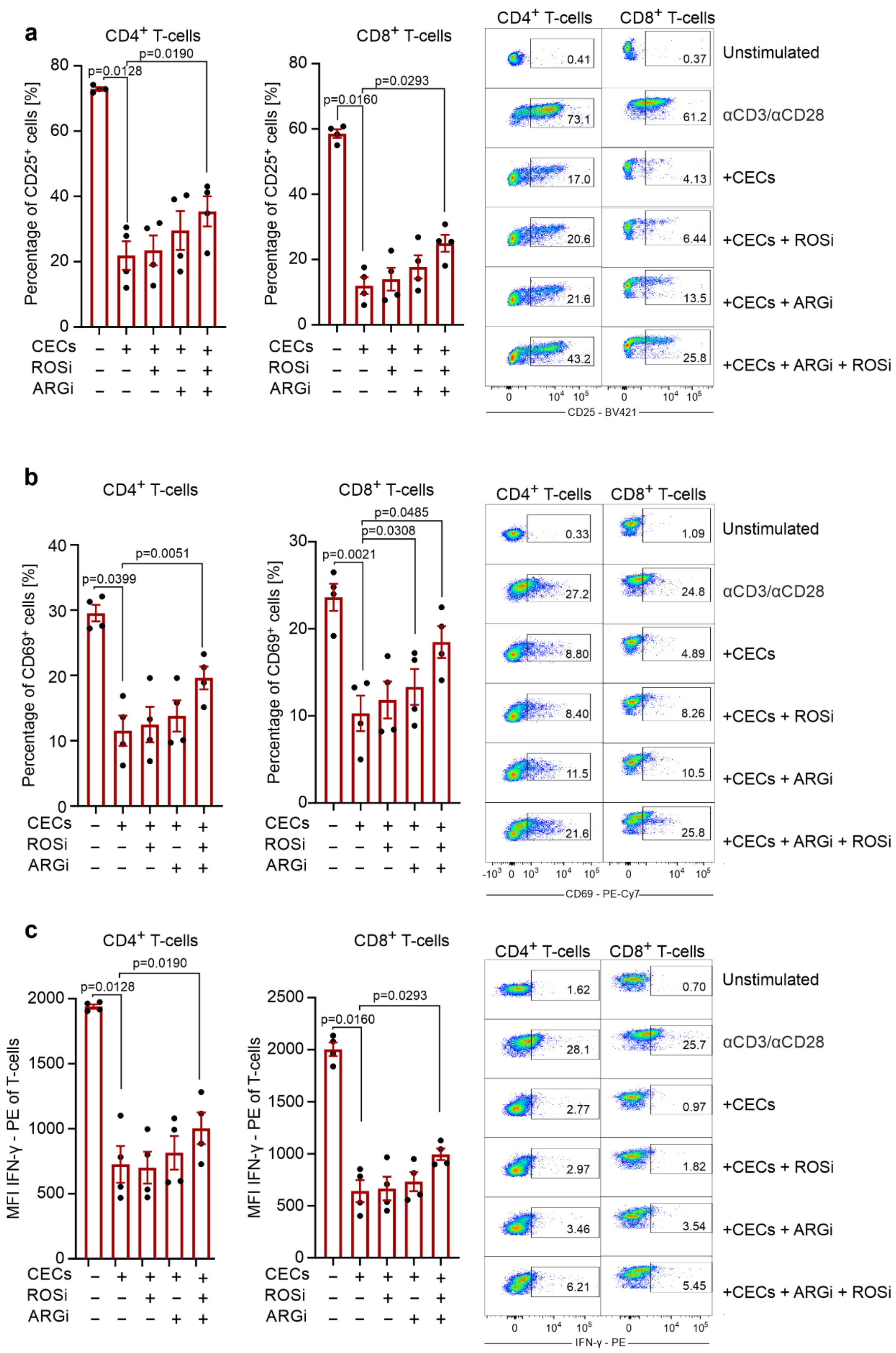

**a,b,** Effects of ARGi (OAT-1746, 1.5 μM) and ROSi (N-acetylcysteine, 200 μM) on $CD4^+$ or $CD8^+$ T-cell activation triggered by αCD3/αCD28 based on the percentage of $CD25^+$ T-cells (**a**) and $CD69^+$ T-cells (**b**). T-cells were co-cultured with PBMC-derived CECs at the stage of $CD71^{high}CD235a^{mid}$ at a ratio of 1:2 for 72h. Representative plots show the fluorescence of CD25 – BV421 (**a**) and CD69 – PE-Cy7 (**b**). **c,** Effects of ARGi (OAT-1746, 1.5 μM) and ROSi (N-acetylcysteine, 200 μM) on interferon γ (IFN-γ) production triggered by αCD3/αCD28 in $CD4^+$ or $CD8^+$ T-cells based on the level of intracellular staining with fluorochrome-labeled antibody. T-cells were co-cultured with PBMC-derived CECs at the stage of $CD71^{high}CD235a^{mid}$ at a 1:2 ratio for 72h. Protein transport inhibitor (BD GolgiStop™) was added for the last 12 hours. Representative plots show the fluorescence of IFN-γ – PE. *P* values were calculated using repeated-measures ANOVA with Sidak's post-hoc tests. Data show means ± SEM. Each point in a, b, c represents data from one individual patient. n values are the numbers of patients used to obtain the data. The source data underlying Supplementary Fig. 22a-c are provided as a Source Data file.

**Supplementary Figure 23. PBMC-derived CECs suppress T-cells proliferation in ARG- and ROS-dependent mechanism**

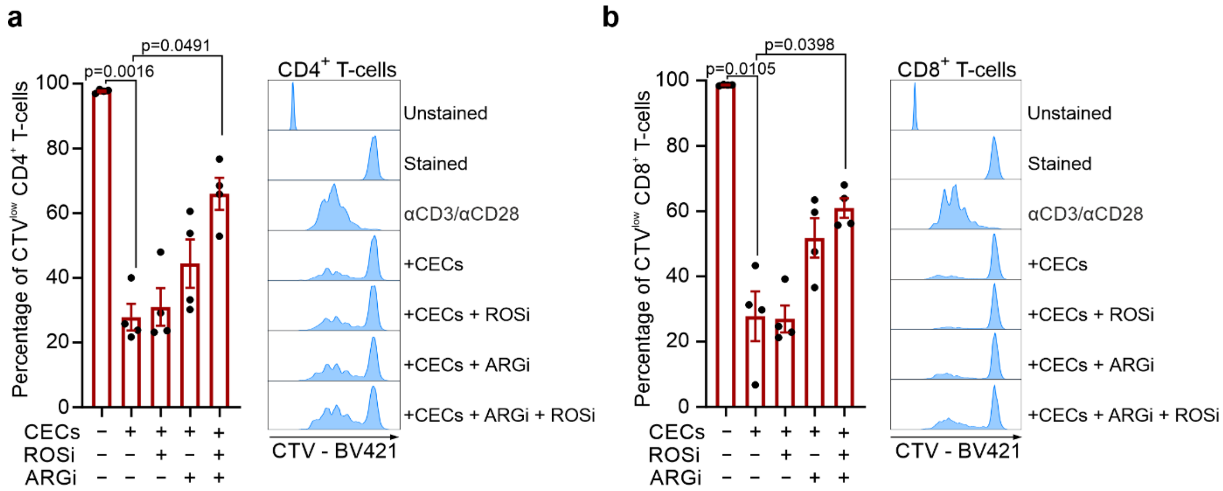


**a,b,** Proliferation triggered by αCD3/αCD28 in CTV-labelled $CD4^+$ (**a,** n=4) and $CD8^+$ (**b,** n=4) T-cells co-cultured with PBMC-derived CECs at ratio 1:2. CECs were at the stage of $CD71^{high}CD235a^{mid}$. *P* values were calculated using repeated-measures ANOVA with Sidak's post-hoc tests. Data show means ± SEM. Each point in a, b represents data from one individual patient. n values are the numbers of patients used to obtain the data. The source data underlying Supplementary Fig. 23a,b are provided as a Source Data file.

**Supplementary Figure 24. Induction of differentiation of K562 cells decreases suppressive effects on T-cells and decreases ARG2 levels**

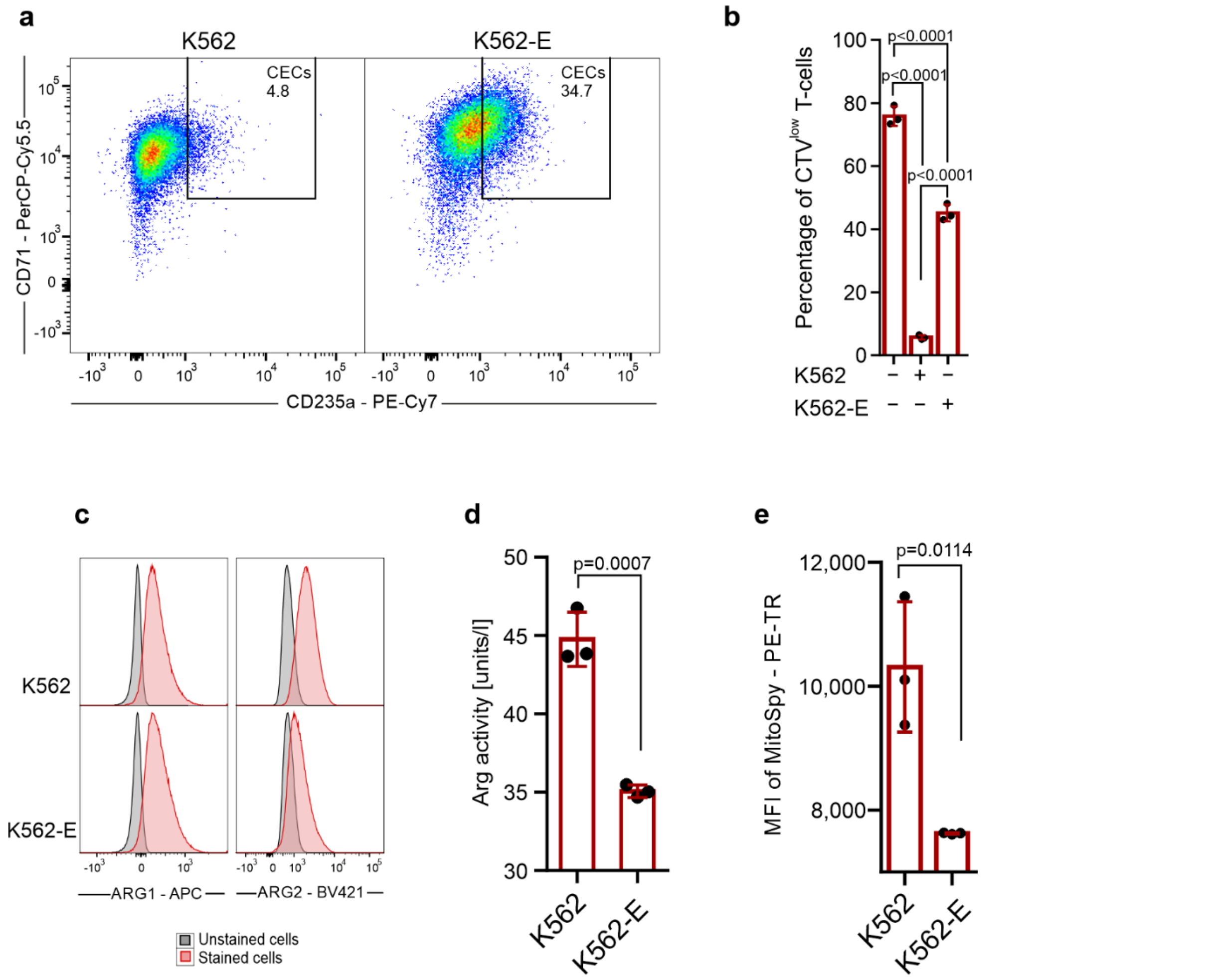


**a,** Representative plots of CD71 and CD235a levels in K562 and sodium butyrate-treated K562 cells (K562-Erythroid, K562-E). **b,** Proliferation of αCD3/αCD28-stimulated CD4$^+$ T-cells in co-culture with K562 and K562-E at a 1:2 ratio. Data show one representative experiment out of two. *P*-values were calculated with one-way ANOVA with Bonferroni's post-hoc test. **c,** Representative histograms of ARG1 and ARG2 expression in erythroid cell lines. **d,** Arginase activity of K562 and K562-E cells calculated per 1×10$^6$ cells (n=3). *P*-value was calculated with an unpaired *t*-test. **e,** Mean Fluorescence Intensity (MFI) of PE-TexasRed-MitoSpy, Mitochondrion probe of K562 and K562-E cells (n=3). *P*-value was calculated with an unpaired *t*-test. Data show means ± SD. Each point in d, e represents data from one biological replicate. n values are the numbers of biological replicates used to obtain the data. The source data underlying Supplementary Fig. 24b, 24d-e are provided as a Source Data file

**Supplementary Figure 25. Erythrocytes have no impact on T-cells proliferation**

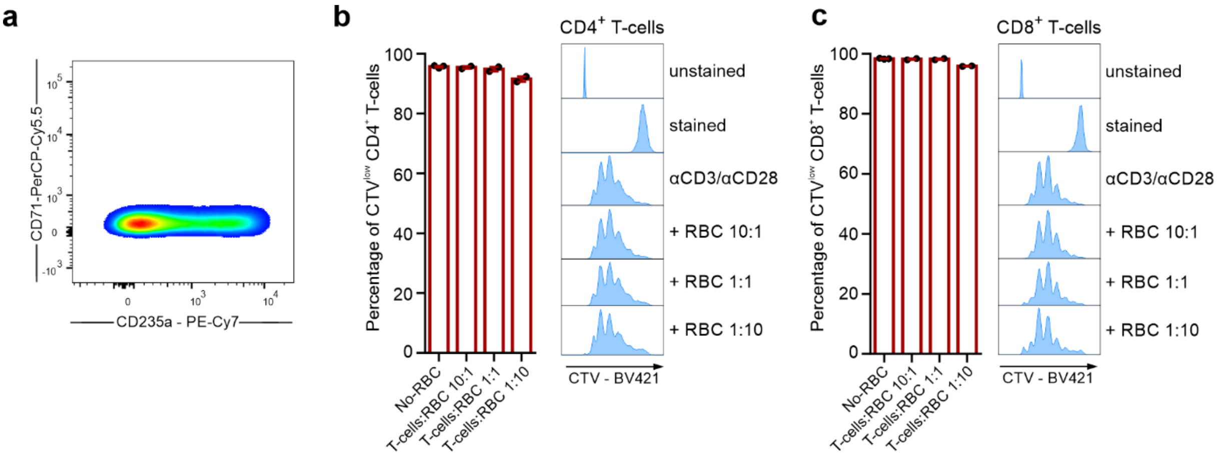


**a,** Representative plot of CD71 and CD235a expression in isolated erythrocytes. **B,c,** Proliferation triggered by αCD3/αCD28 in CTV-labelled $CD4^+$ (**b)** and $CD8^+$ (**c)** T-cells co-cultured with RBC (n=2). Data show means ± SEM. Each point in b,c represents data from an individual patient. n values are the numbers of patients used to obtain the data. The source data underlying Supplementary Fig. 25b,c are provided as a Source Data file.

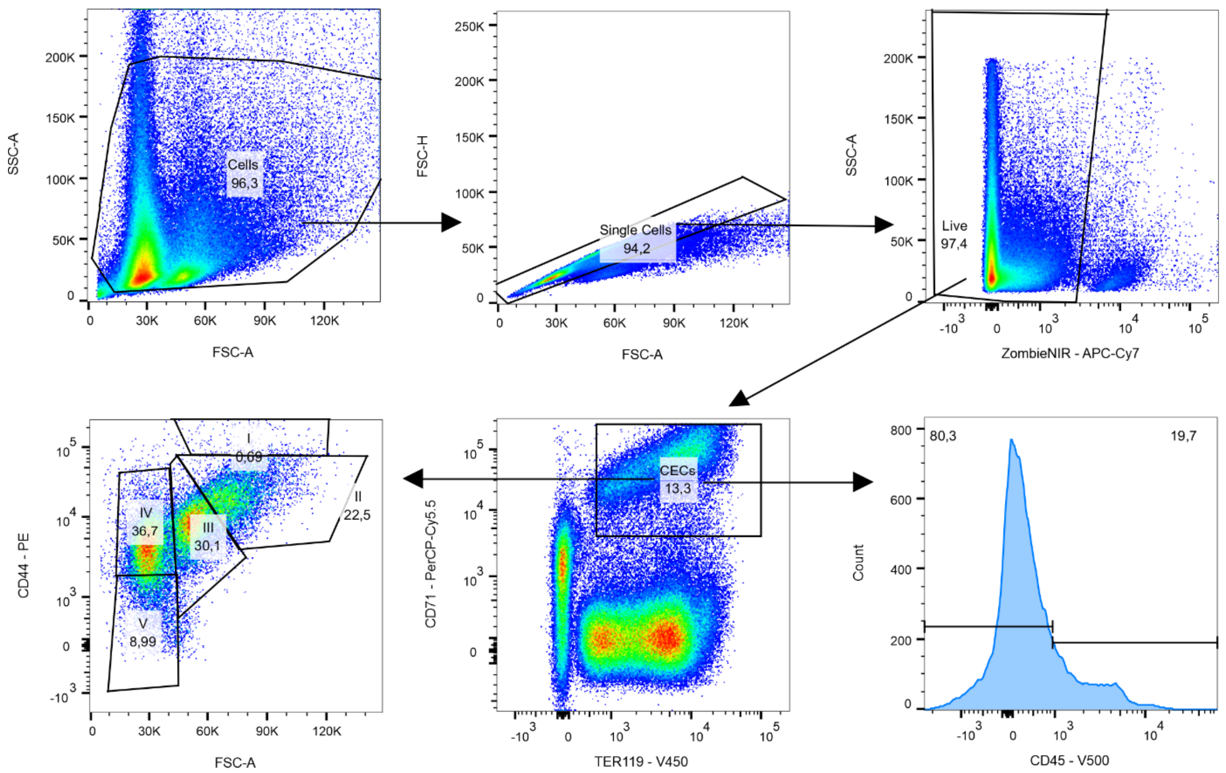


**Supplementary Figure 26**. Gating strategy used to analyze the data shown in Figure 1a-b, 1f-g, Supplementary Figure 2a-c, Supplementary Figure 6a-g, Supplementary Figure 13a. A minimum of 10 000 cells were acquired within the CD45$^+$ CECs gate.

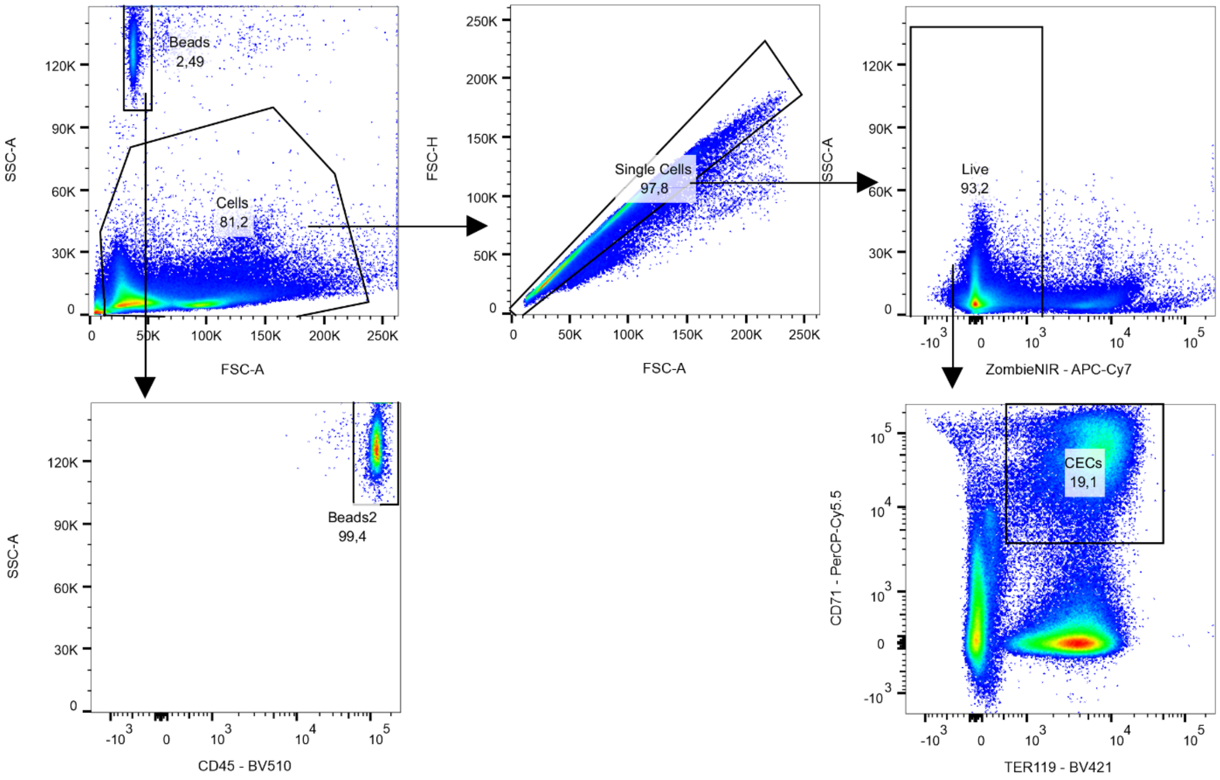


**Supplementary Figure 27.** Gating strategy used to analyze the data shown in Figure 1c. A minimum of 10 000 cells were acquired within the CECs gate.

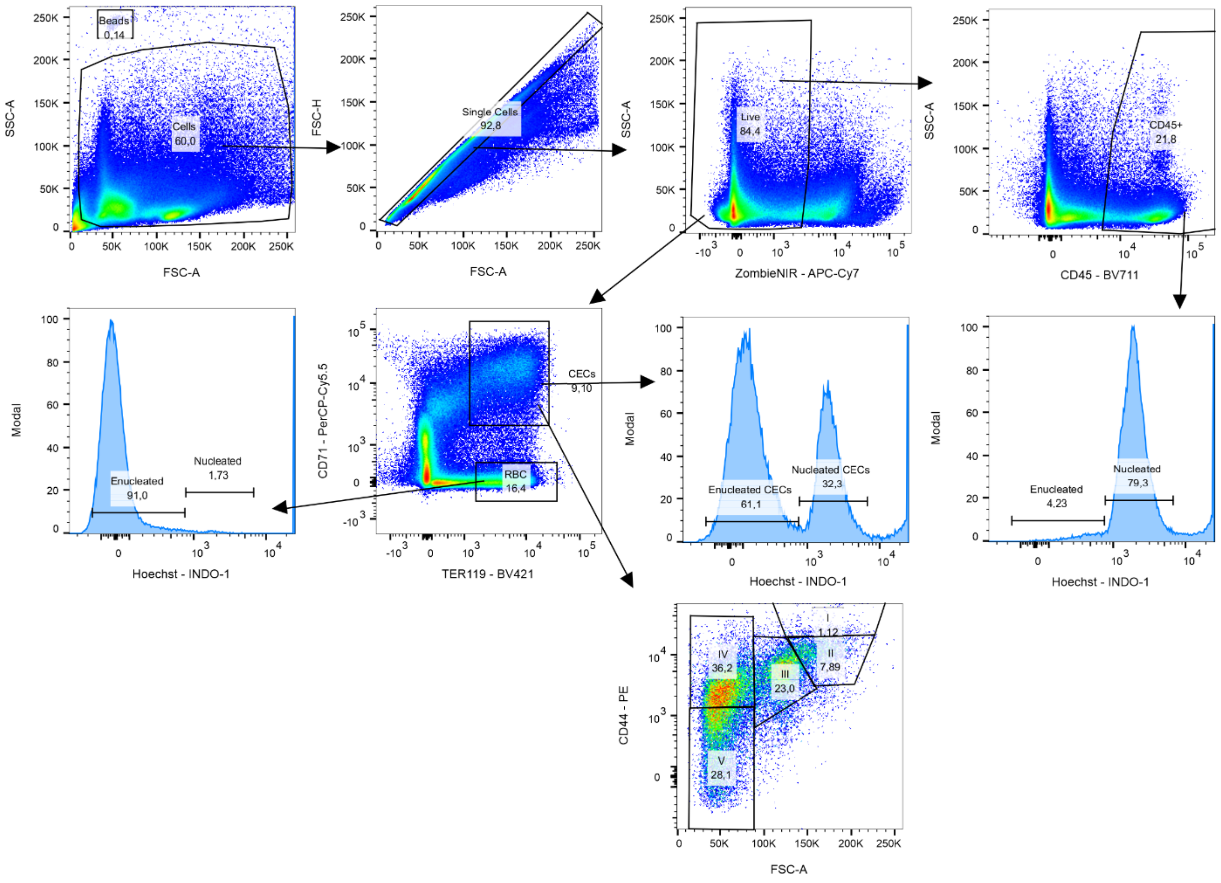


**Supplementary Figure 28.** Gating strategy used to analyze the data shown in Figure 1g-i. A minimum of 30 000 cells were acquired within the CECs gate.

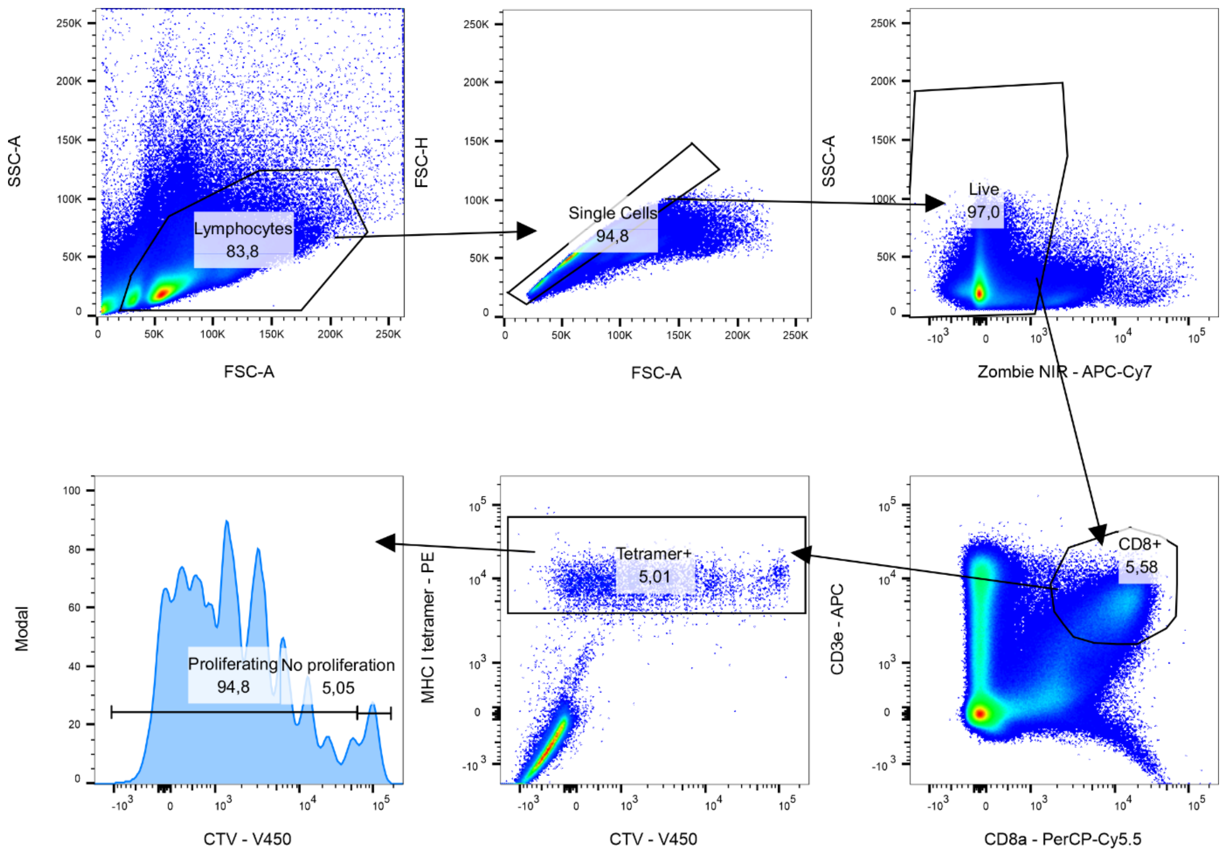


**Supplementary Figure 29.** Gating strategy used to analyze the data shown in Figure 2b. A minimum of 40 000 cells were acquired within the CD8$^+$ cells gate. The gate for proliferating (CTV$^{low}$) Tetramer$^+$ CD8a$^+$ T-cells was set based on the unstimulated control. Proliferation histograms were generated in FlowJo v10.6.2.

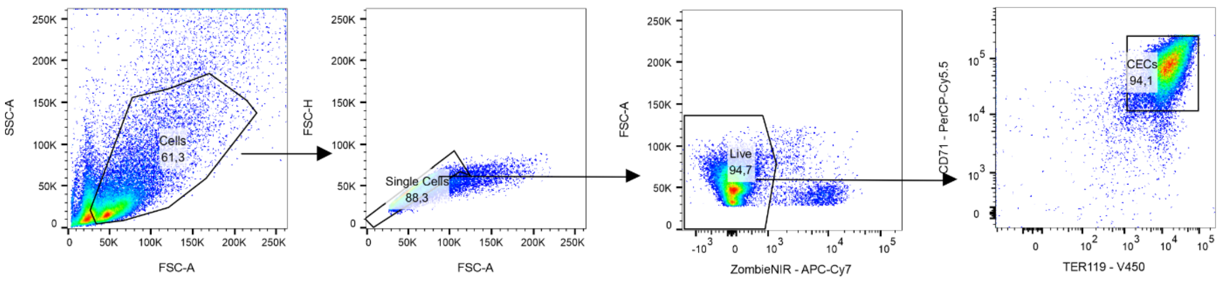


**Supplementary Figure 30.** Gating strategy used to analyze the data shown in Figure 2c, Supplementary Figure 4a. A minimum of 20 000 cells were acquired within the live cells gate.

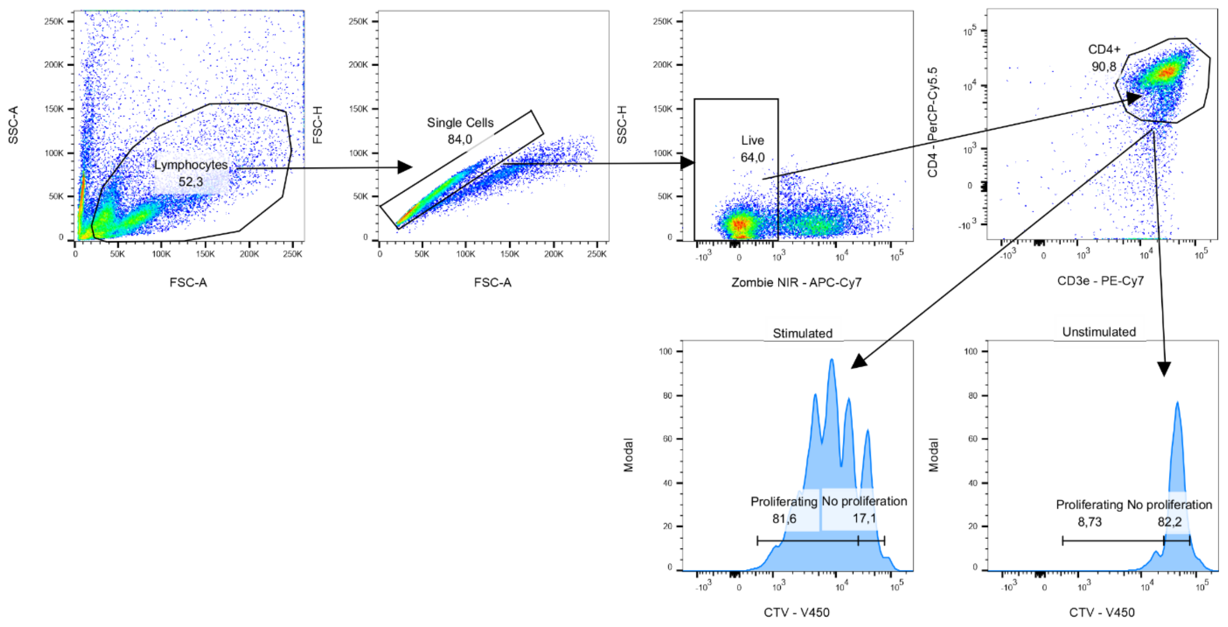


**Supplementary Figure 31.** Gating strategy used to analyze the data shown in Figure 2d, Figure 5b-f, Figure 6c, Figure 8e-f, Figure 9b-I, Figure 10b-c, Figure 10e-g, Supplementary Figure 8a-b, Supplementary Figure 11a, Supplementary Figure 15c-d, Supplementary Figure 18a-b, Supplementary Figure 21d-e, Supplementary Figure 23a-b, Supplementary Figure 24b, Supplementary Figure 25b-c. Exemplary gating strategy used for the murine CD4$^{+}$ T-cell population (analogous strategy was used for the human CD4$^{+}$ T-cell population and murine and human CD8$^{+}$ T-cell population). A minimum of 10 000 cells were acquired within the lymphocytes gate. The gate for proliferating (CTV$^{low}$) CD4$^{+}$ T-cells was set based on the unstimulated control. Proliferation histograms were generated in FlowJo v10.6.2.

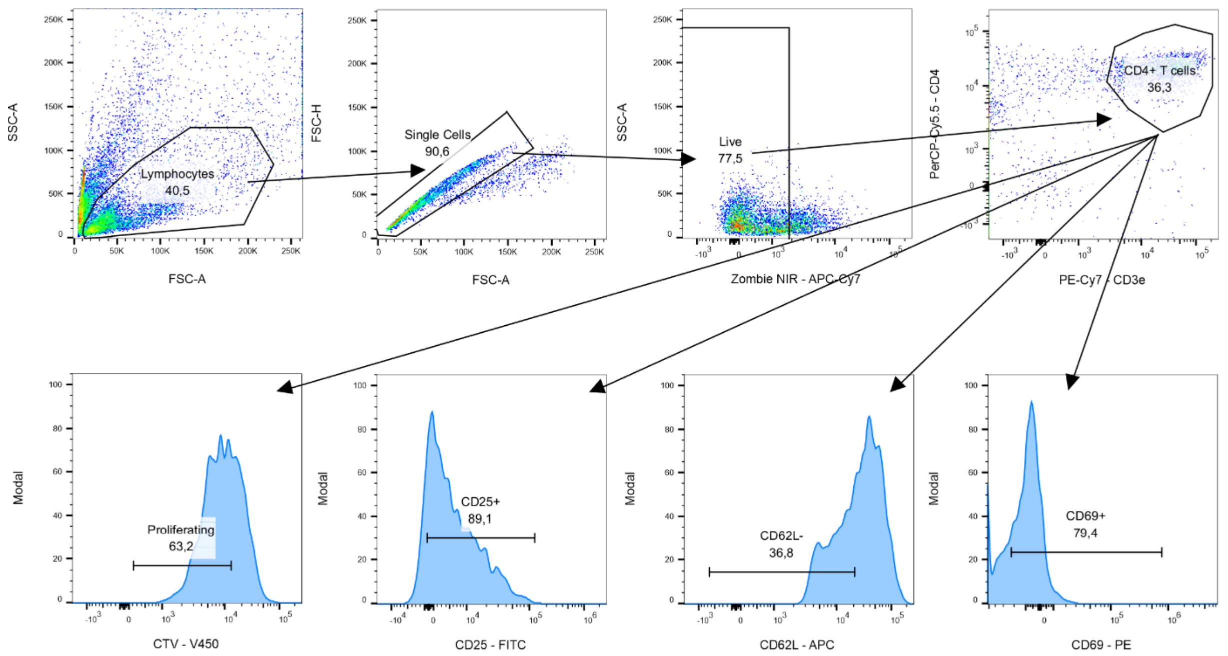


**Supplementary Figure 31.** Gating strategy used to analyze the data shown in Figure 5a. A minimum of 10 000 cells were acquired within the lymphocytes gate. The gate for proliferating ($CTV^{low}$) cells as well as $CD25^{+}$, $CD62L^{-}$ and $CD69^{+}$ cells was set based on the unstimulated control. Proliferation histograms were generated in FlowJo v10.6.2.

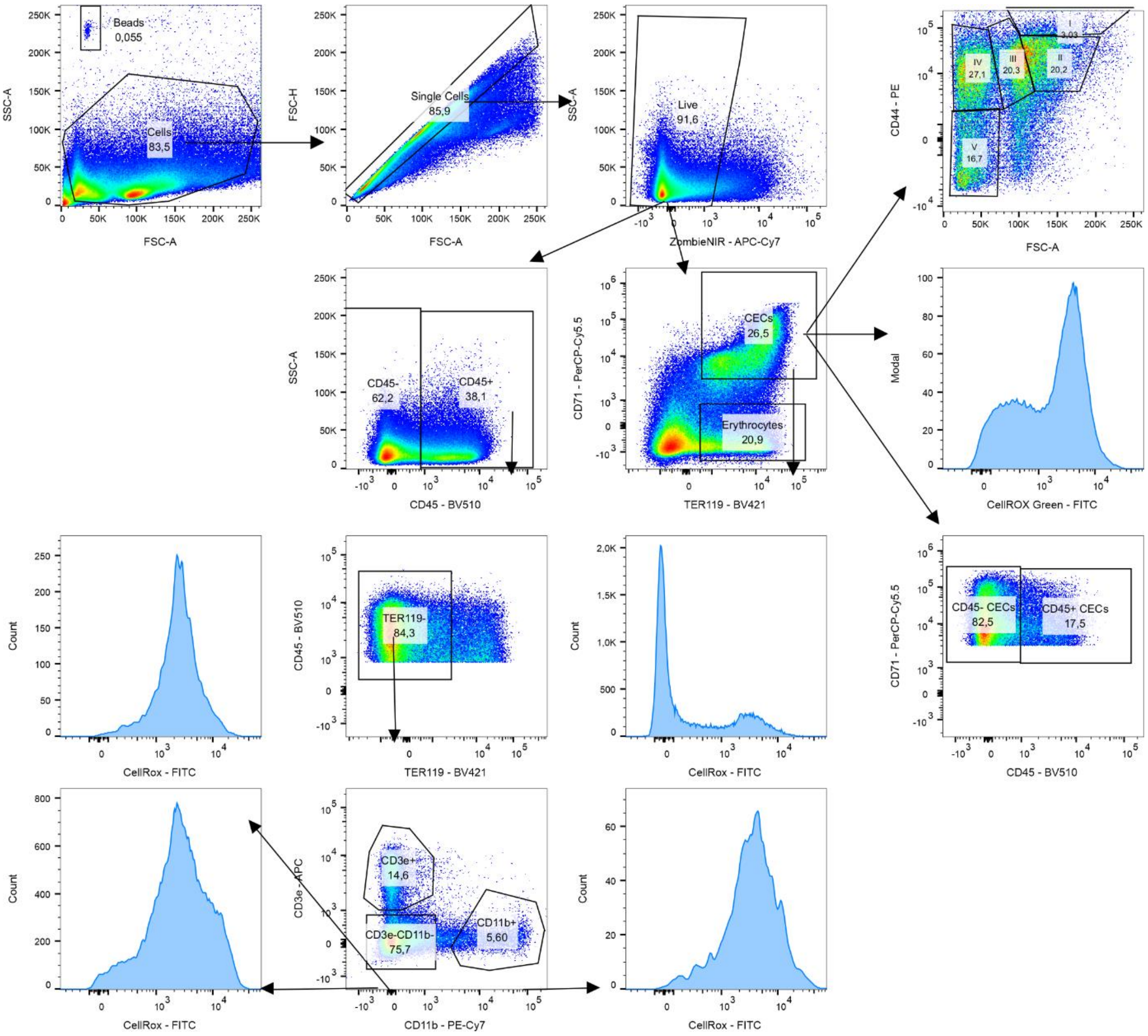


**Supplementary Figure 32.** Gating strategy used to analyze the data shown in Figure 3a, Figure 3b, Supplementary Figure 5a-f. A minimum of 100 000 cells were acquired within the live cell gate. Histograms were generated in FlowJo v10.6.2.

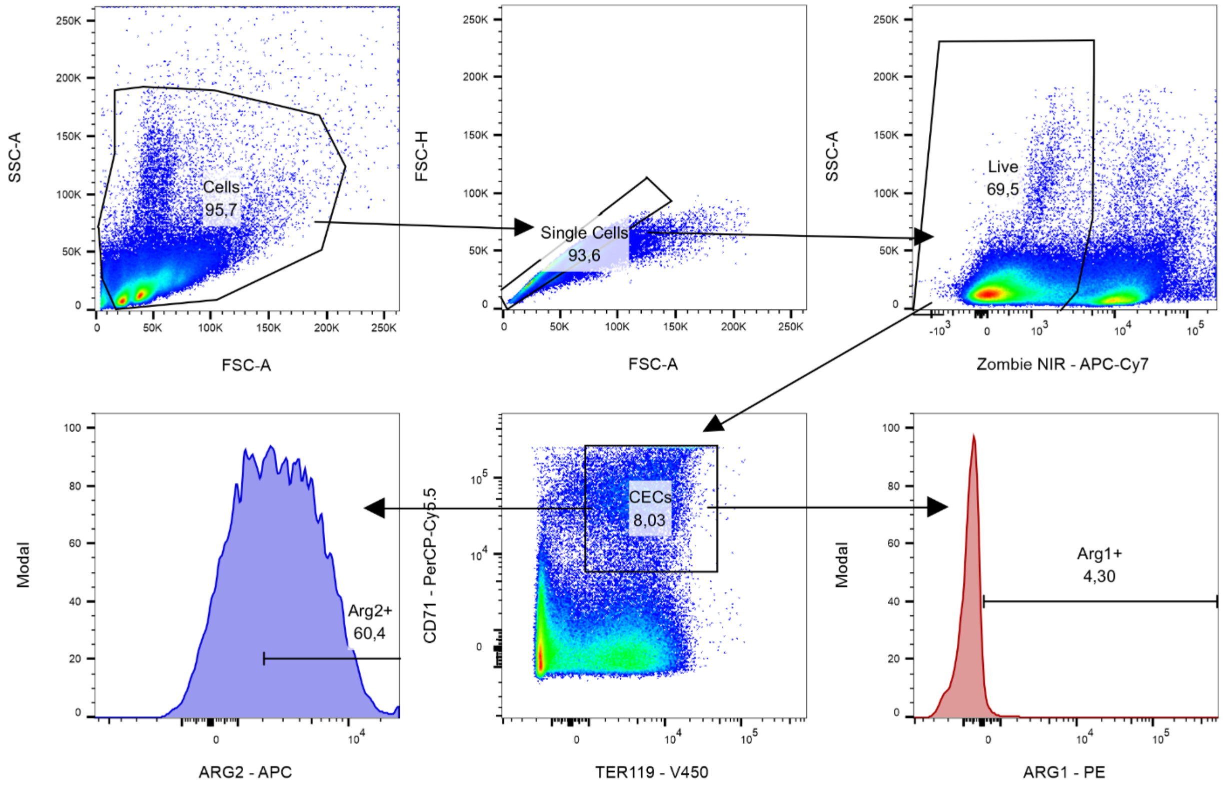


**Supplementary Figure 33.** Gating strategy used to analyze the data shown in Figure 3c-e, Supplementary Figure 14b-c. A minimum of 20 000 cells were acquired within the CECs cells gate. The gate for ARG1$^{+}$ cells and ARG2$^{+}$ was set based on the Fluorescence Minus One (FMO) control. Histograms were generated in FlowJo v10.6.2.

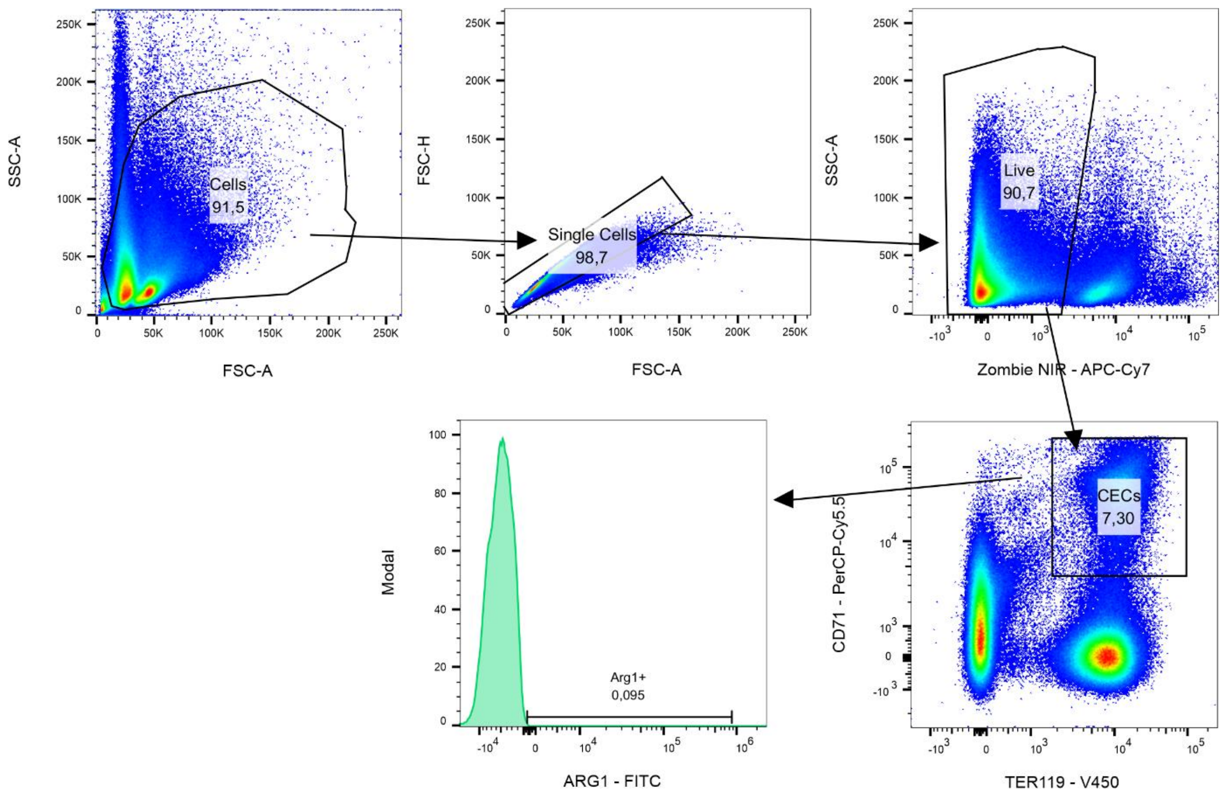


**Supplementary Figure 34.** Gating strategy used to analyze the data shown in Figure 3f-h, Figure 3k-l, Supplementary Fig. 10a-d. A minimum of 20 000 cells were acquired within the CECs cells gate. The gate for YFP/ARG1$^+$ cells was set based YFP negative control from C57BL/6 mice. Histograms were generated in FlowJo v10.6.2.

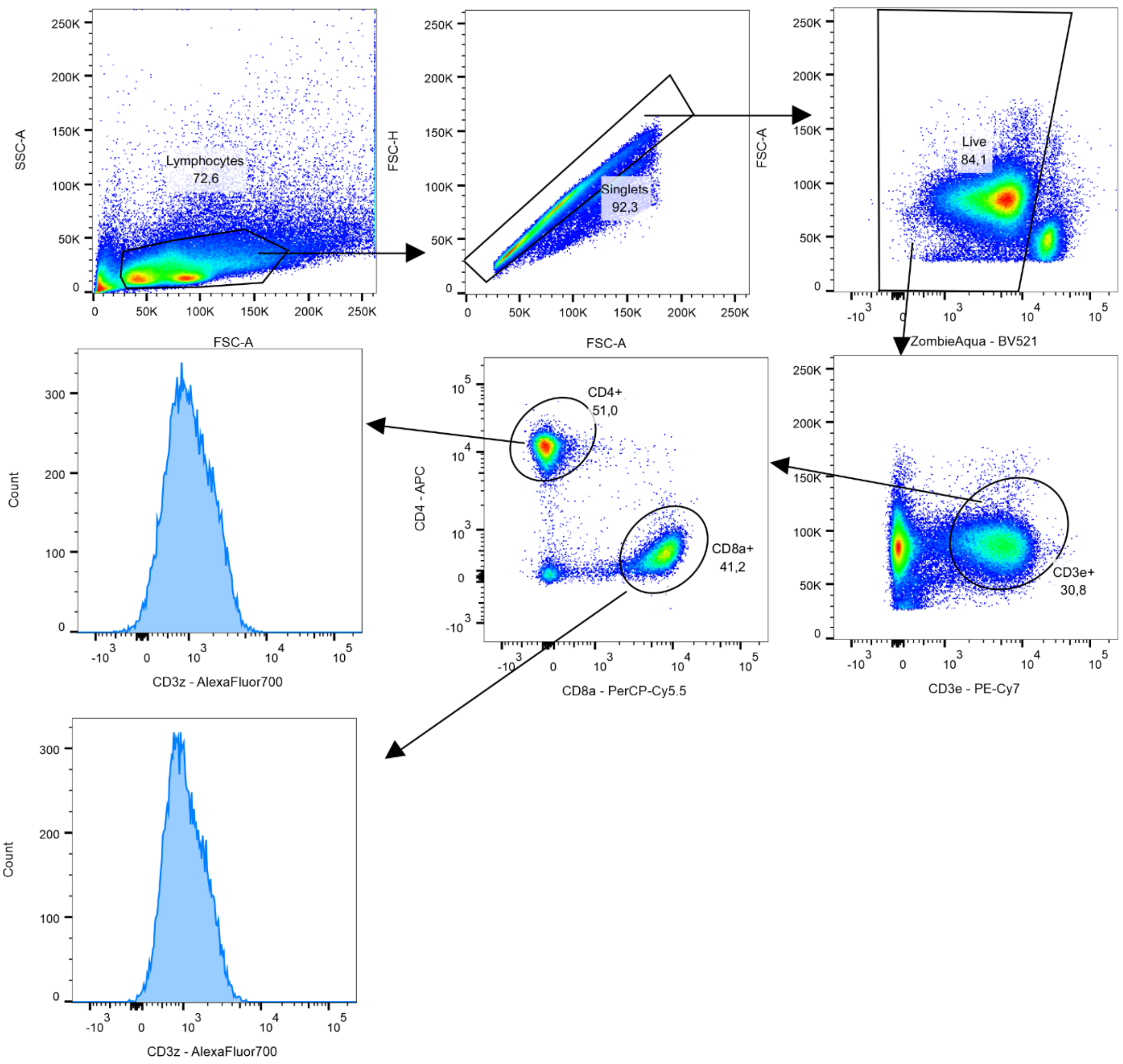


**Supplementary Figure 35.** Gating strategy used to analyze the data shown in Figure 5k-l, Supplementary Figure 13b-c. A minimum of 100 000 cells were acquired within the live cells gate. Histograms were generated in FlowJo v10.6.2.

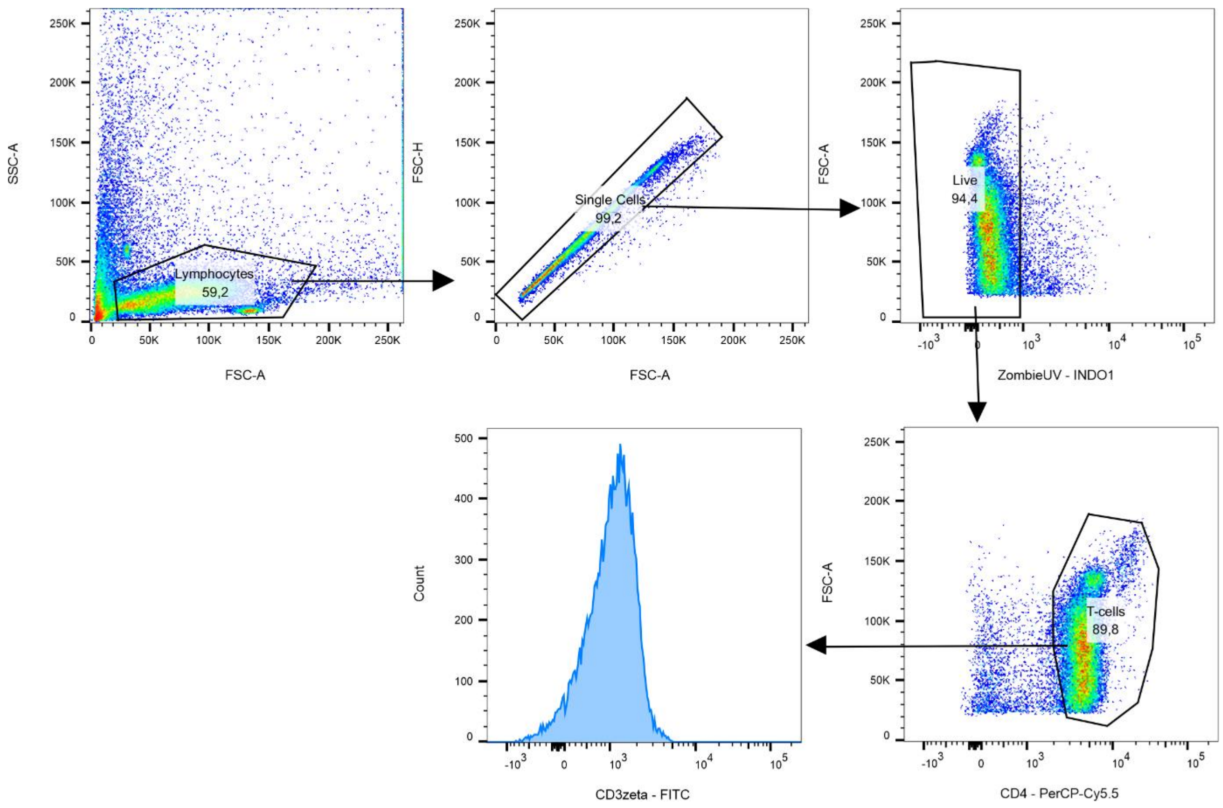


**Supplementary Figure 36.** Gating strategy used to analyze the data shown in Figure 5m-n, Figure 6a-b,. A minimum of 10 000 cells were acquired within the T-cells gate. Exemplary gating strategy used for the CD4$^+$ T-cell population (analogous strategy was used for the murine CD8$^+$ T-cell population). Histograms were generated in FlowJo v10.6.2.

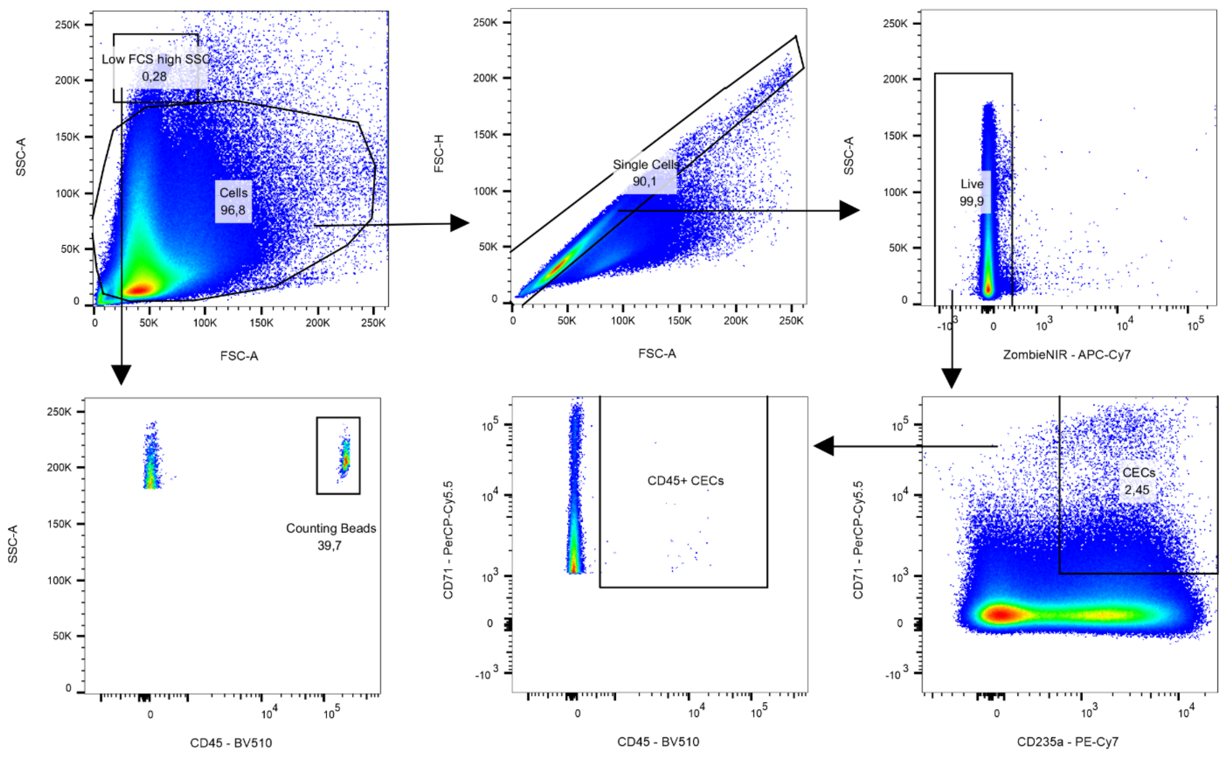


**Supplementary Figure 37.** Gating strategy used to analyze the data shown in Figure 7a-g, Figure 8a, Supplementary Figure 15a-b, Supplementary Figure 16g, Supplementary Figure 22c, Supplementary Figure 25a. A minimum of 500 000 cells were acquired within the live cells. For the data shown in Figure 7f-g, Figure 8a, Supplementary Figure 22a a minimum of 50 000 were acquired within the live cells.

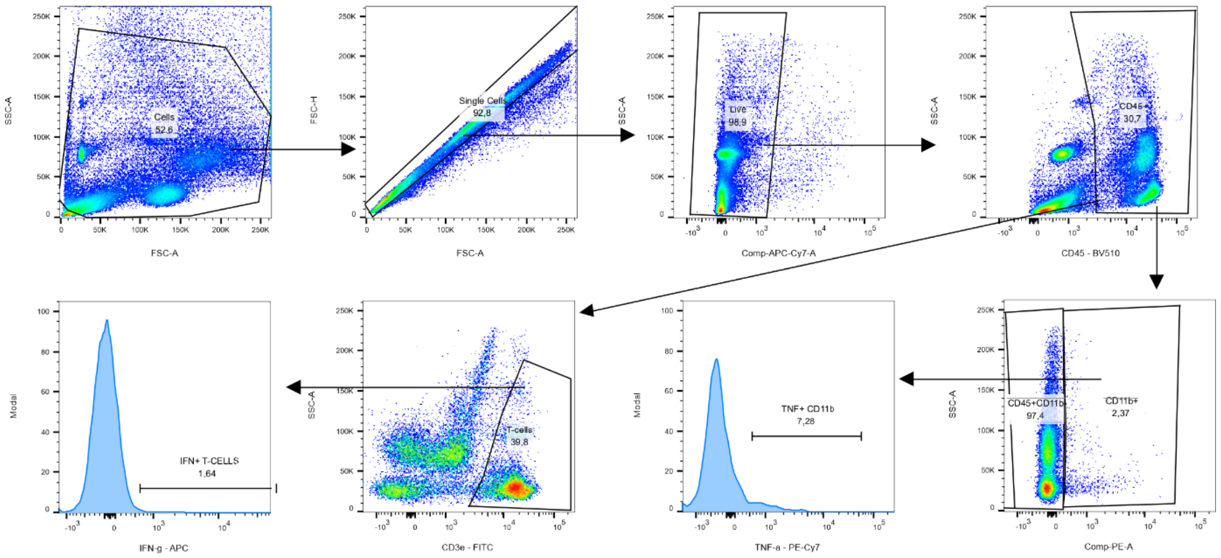


**Supplementary Figure 38.** Gating strategy used to analyze the data shown in Figure 7h-i, Supplementary Fig. 15e. A minimum of 100 000 cells were acquired within the live cells. The gate for IFN-γ$^+$ and TNF-α$^+$ cells were set based on unstimulated controls.

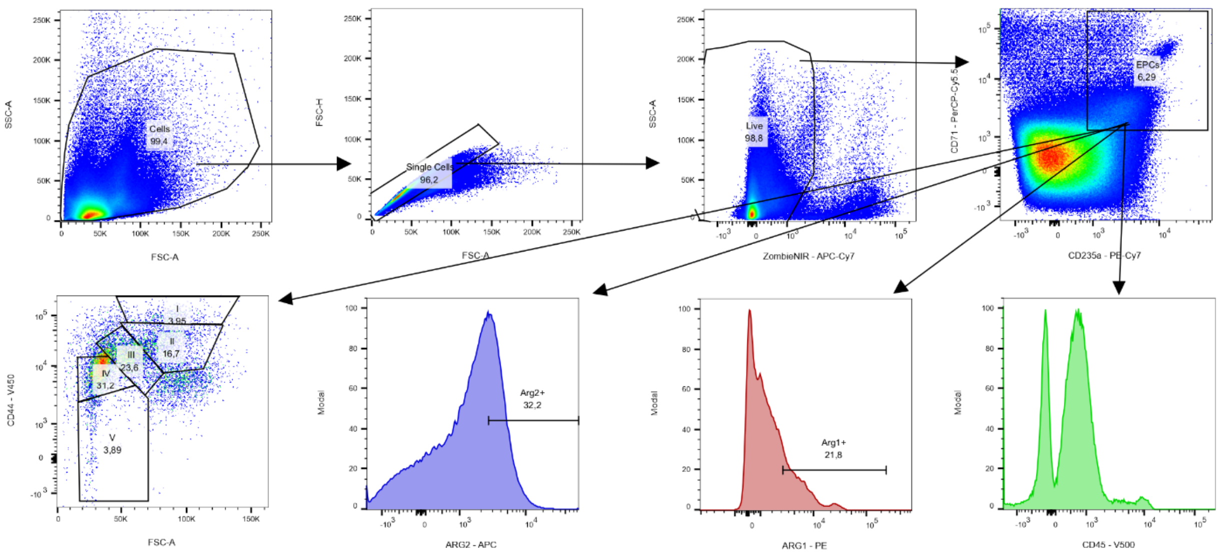


**Supplementary Figure 39.** Gating strategy used to analyze the data shown in Figure 8b-d, Supplementary Fig. 16a-f. A minimum of 100 000 cells were acquired within the live cells. The gate for ARG1$^{+}$ and ARG2$^{+}$ cells were set based on fluorescence minus one (FMO) controls.

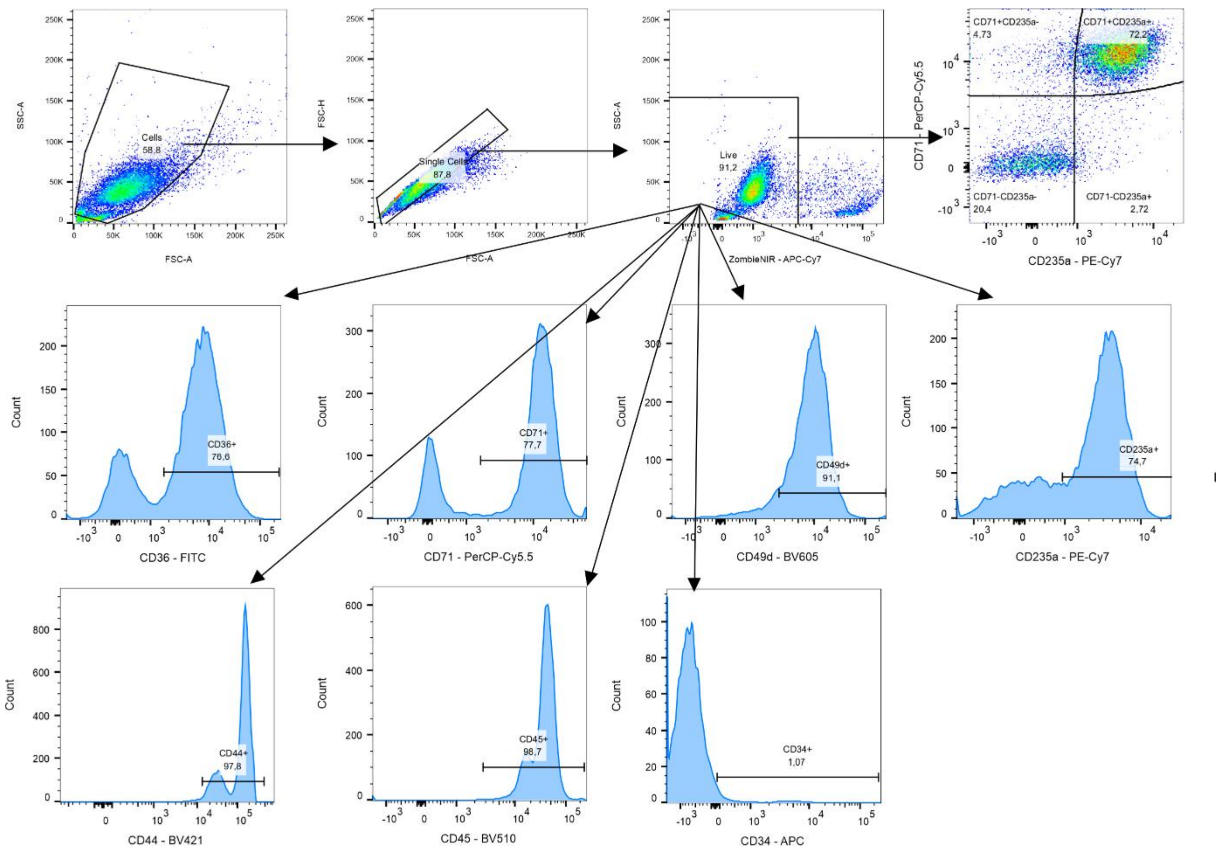


**Supplementary Figure 40.** Gating strategy used to analyze the data shown in Figure 9a, Figure 10a, Figure 10d, Figure 10h-j, Supplementary Fig. 19a,b,. A minimum of 50 000 cells were acquired within the live cells. The gate for CD71$^+$, CD235a$^+$, CD36$^+$, CD49d$^+$, CD44$^+$, CD45$^+$, CD34$^+$ cells were set based on fluorescence minus one (FMO) controls.

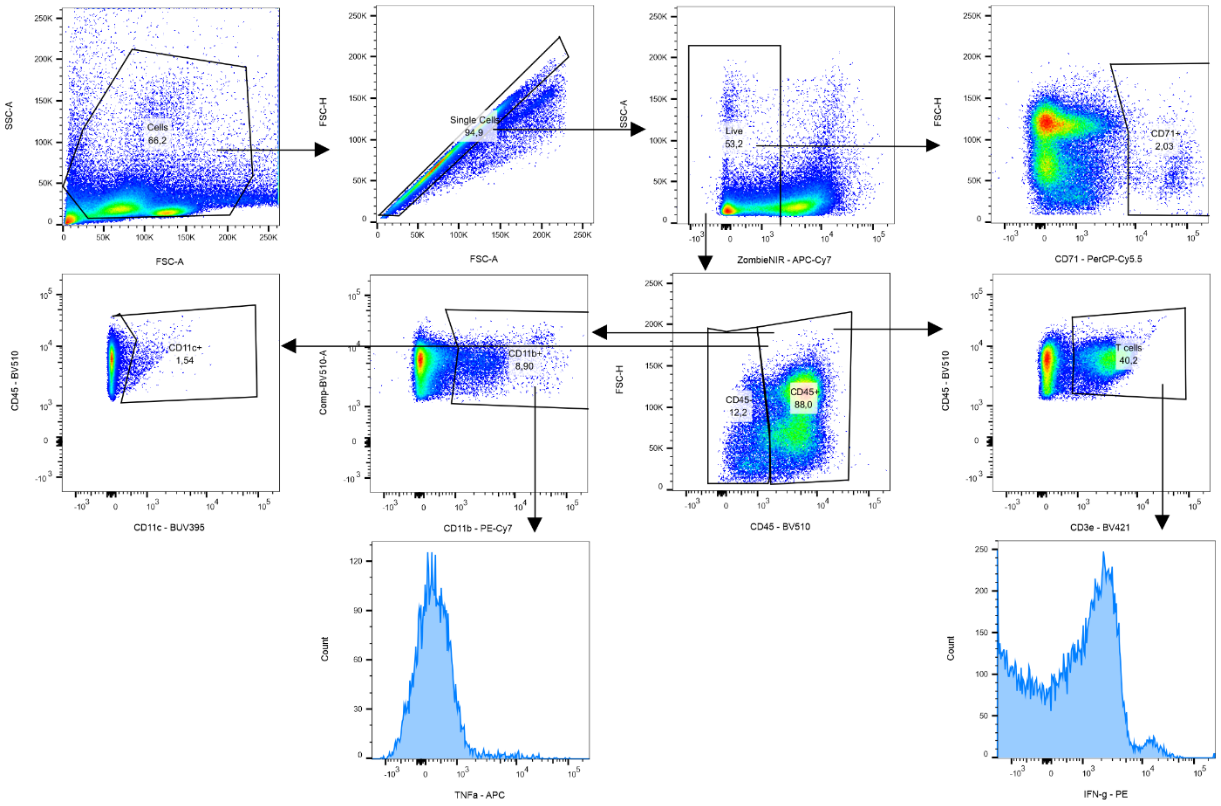


**Supplementary Figure 41.** Gating strategy used to analyze the data shown in Supplementary Figure 3a-b, Supplementary Figure 3d. A minimum of 50 000 cells were acquired within the live cells. The gate for TNF-α$^+$ and IFN-γ$^+$ cells were set based on isotype controls.

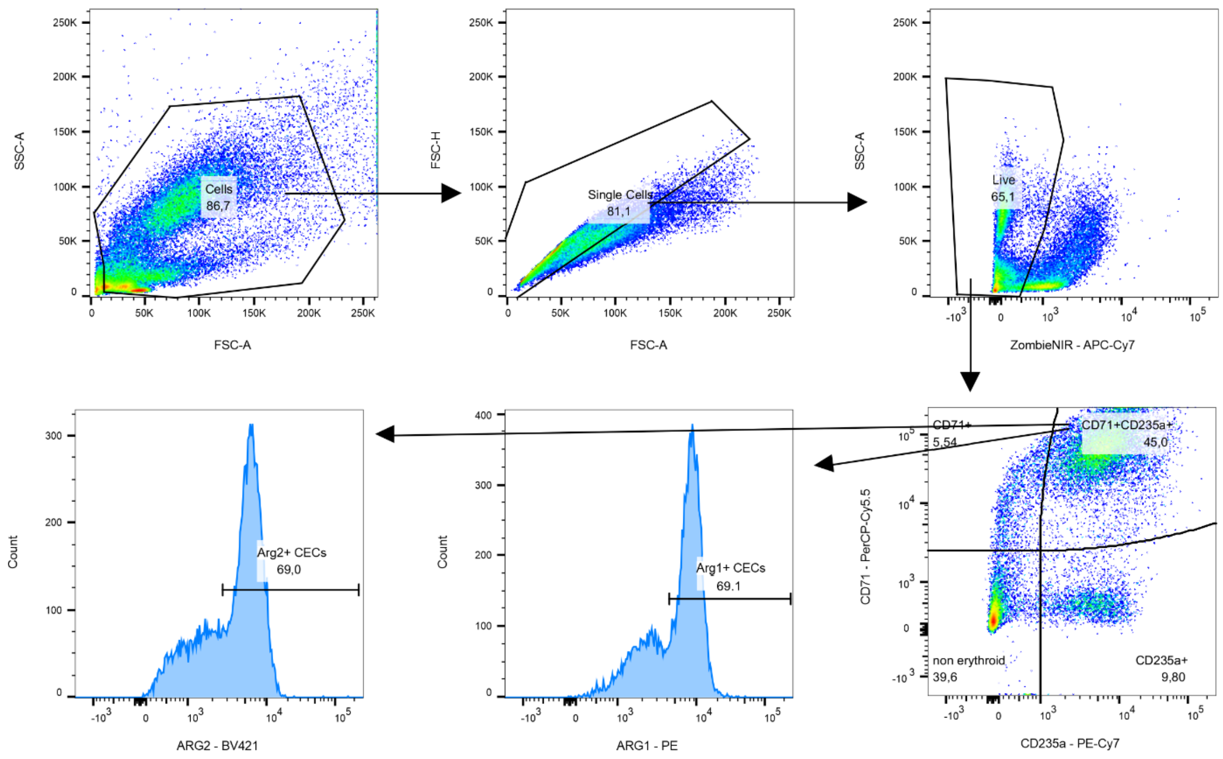


**Supplementary Figure 42.** Gating strategy used to analyze the data shown in Supplementary Figure 17b-d, Supplementary Figure 20a-b, Supplementary Figure 24a, Supplementary Figure 24c. A minimum of 20 000 cells were acquired within the live cells. The gate for ARG1$^+$ and ARG2$^+$ cells were set based on fluorescence minus one (FMO) controls.

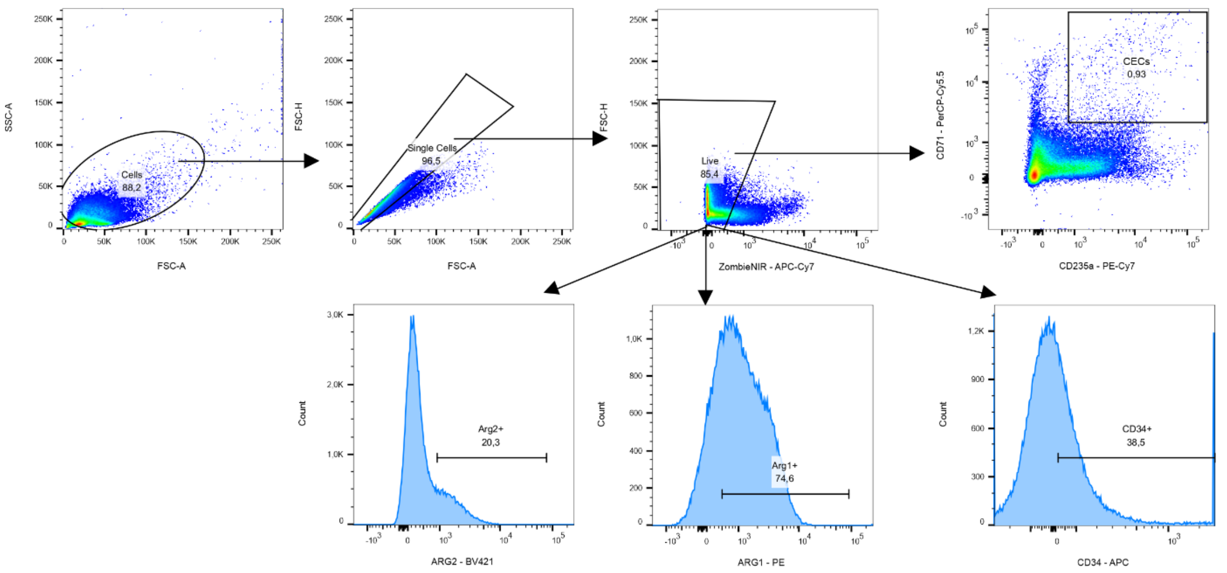


**Supplementary Figure 43.** Gating strategy used to analyze the data shown in Supplementary Figure 21a-c. A minimum of 50 000 cells were acquired within the live cells. The gate for $CD34^+$, $ARG1^+$ and $ARG2^+$ cells were set based on fluorescence minus one (FMO) controls.

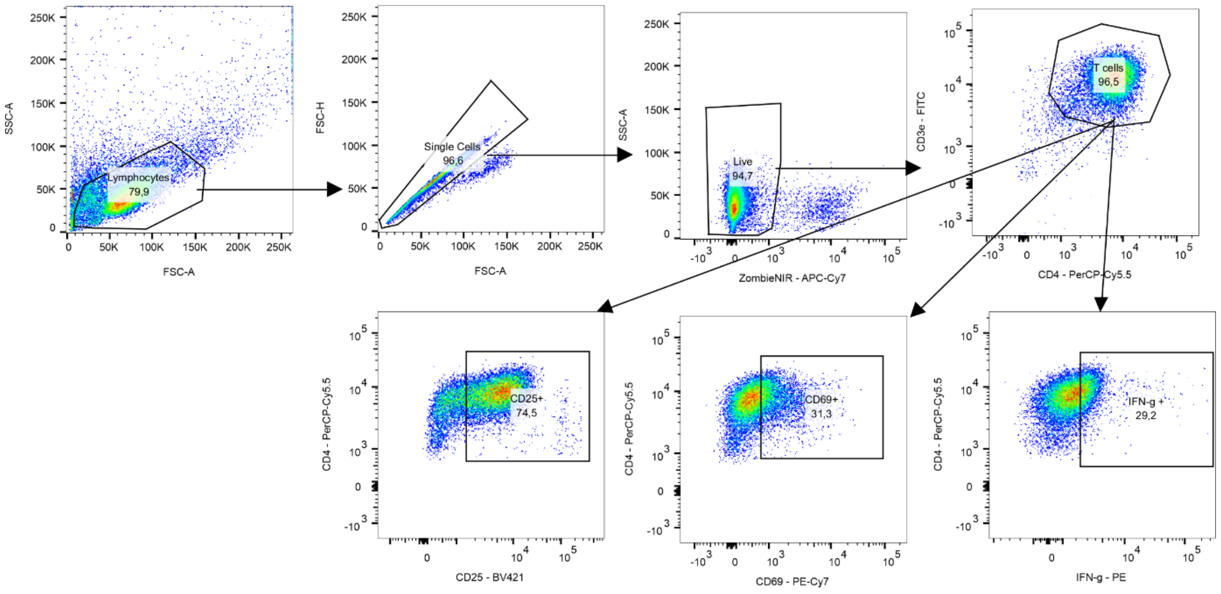


**Supplementary Figure 44.** Gating strategy used to analyze the data shown in Supplementary Figure 22a-c. A minimum of 20 000 cells were acquired within the live cells. The gate for CD69$^+$ and CD25$^+$ were set based on the unstimulated control. The gate for IFN-γ$^+$ cells were set based on the isotype control.
